# PRC2 complex subunit JARID2 regulates embryonic pituitary stem cell differentiation to the POMC lineage

**DOI:** 10.64898/2026.06.11.731685

**Authors:** Zihuan Lin, Michelle L. Brinkmeier, Karen E. Weis, Hayley Bossard, Lori T. Raetzman

## Abstract

The pituitary is essential for growth, fertility, and stress response. Pituitary development is defined by transcription factor cascades, but how chromatin landscapes influence it remains elusive. JARID2 is a subunit of polycomb repressive complex 2 repressing transcription by depositing trimethylation on histone 3 lysine 27 (H3K27me3). Here, we established that H3K27me3 levels increase with differentiation in anterior pituitary from embryonic day (E) 10.5 to E14.5 despite ubiquitous *Jarid2* mRNA expression. Germline loss of *Jarid2* causes pituitary stem cell (PSC) hyperplasia and corticotrope hypoplasia without changing H3K27me3 bulk levels at E14.5. Pituitary-specific loss of *Jarid2* with *Prop1^T2AiCre^* (*Jarid2^Pitko/Pitko^*) exhibits POMC lineage hypoplasia and PSC hyperplasia at E16.5. *Pou1f1* or *Nr5a1* lineages are not affected by *Jarid2* mutation. To determine gene expression and chromatin accessibility changes in *Jarid2^Pitko/Pitko^*, we performed single nucleus (sn) multi-omics on E14.5 pituitaries. SnRNAseq revealed potential targets of *Jarid2*, including *Meis2*. SnATACseq revealed chromatin reprograming towards PSC retention rather than differentiation. *Pax7* is reduced in both modalities in PSC. Together, we show that *Jarid2* promotes stemness exit and POMC lineage differentiation, uncovering novel epigenetic regulation of pituitary development.

**SUMMARY STATEMENT:** Expanding the repertoire of the regulation of pituitary development, chromatin modifier also help stem cells in this essential gland to differentiate and choose hormone-secreting fate.

## INTRODUCTION

Regarded as the “master gland”, the pituitary gland is crucial for the regulation of reproduction, growth, metabolism, and the stress response. To orchestrate these physiological processes, mouse pituitaries differentiate into 3 lobes responsible for different roles. The posterior lobe (PL) serves as a signaling hub for development of the oral ectoderm into anterior pituitary, along with the ventral diencephalon. In adulthood, the PL contains specialized glia cells (pituicytes) and axon terminals of arginine vasopressin (AVP) and oxytocin (OXT) producing neurons (Drouin and Brière, 2022). The anterior lobe (AL) consists of 5 hormone-secreting cell types: corticotropes (adrenocorticotropin, ACTH), somatotropes (growth hormone, GH), thyrotropes (thyroid-stimulating hormone, TSH), lactotropes (prolactin, PRL), and gonadotropes (luteinizing hormone, LH and follicle-stimulating hormone, FSH) (Davis et al., 2013). Additionally, melanotropes in the intermediate lobe (IL) are responsible for secreting melanocyte stimulating hormone (αMSH). Because corticotropes and melanotropes both contain pro-opiomelanocortin (POMC), they are also regarded as POMC lineage (Drouin and Brière, 2022).

If any of these cell types fail to form during gestation, the pituitary gland will have hormonal deficiencies, resulting in congenital hypopituitarism (CH). Currently, known CH related genetic variants are mainly transcription factors (TFs) and signaling factors important for pituitary development including *Prop1* and *Pou1f1*, but 84% of the genetic etiology of CH remains uncharacterized (Gregory and Dattani, 2020). To understand the genetic pathology of CH, we need to better understand pituitary development.

Pituitary development starts with pituitary stem cells marked with *Sox2* and *Prop1* (Davis et al., 2016a; Fauquier et al., 2008). Endocrine cellular differentiation is regulated by a TF cascade (reviewed in Edwards et al. (2016)). *Pitx1/2* and *Lhx3* determine pituitary identity from oral ectoderm (Gregory and Dattani, 2020). During differentiation, hormone secreting cell types fall into 3 lineages categorized by lineage-specific TFs: *Tbx19* determines the POMC lineage; *Pou1f1* lineage includes thyrotropes, somatotropes and lactotropes; *Nr5a1* lineage develops into gonadotropes (Drouin and Brière, 2022).

Therefore, the fidelity of pituitary gene expression in each lineage is vital for proper pituitary development. One understudied mechanism of pituitary gene expression fidelity maintenance is the regulation on the higher-order structure of chromatin, namely epigenetics. Previous studies have underscored the importance of epigenetics in pituitary differentiation. For example, *Insm1* can regulate differentiation of all pituitary lineages by recruiting histone modifiers (Welcker et al., 2013). Among the three lineages, multiple studies have shown the epigenetic regulation in NR5A1 lineage (Laverrière et al., 2016; Pacini et al., 2019; Xie et al., 2017). And the POMC lineage is well-defined for *Pax7*-mediated epigenetic regulation of the differentiation towards melanotrope instead of corticotrope fate (Budry et al., 2012; Gouhier et al., 2024).

Using mouse models from the International Mouse Phenotyping Consortium (IMPC), we identified that loss of function (LOF) in epigenetic factor *Jarid2* causes anterior lobe malformation and aberrant Rathke’s pouch patterning (Martinez-Mayer et al., 2024a). JARID2 is a subunit of Polycomb Repressive Complex 2 (PRC2) specific to variant PRC2.2. The PRC2 complex is responsible for adding trimethylation of histone 3 at lysine 27 (H3K27me3) to silence gene expression (Loh and Veenstra, 2022). *Jarid2* is important for both recruiting PRC2 to the genome (Kasinath et al., 2021) and limiting the over expansion of H3K27me3 (Agius et al., 2026).

*Jarid2* is important for stem cell differentiation in many systems. In early embryonic development for example, *Jarid2* suppresses the ectodermal fate and the formation of early differentiating intermediates during exit of pluripotency (Loh et al., 2021). Conversely, human mutations in *JARID2* have been associated with craniofacial malformation (Hao et al., 2015), developmental delay, and neurodevelopmental disorders (Verberne et al., 2021). These phenotypes suggest that *Jarid2* may contribute to CH pathogenesis. However, the function of *Jarid2* in pituitary development remains unknown.

Here, we show that despite the ubiquitous expression pattern of PRC2 subunits, H3K27me3 levels increase with stem cell differentiation in embryonic pituitary development. Germline loss of *Jarid2* results in excess pituitary stem cells and failure to differentiate corticotrope cells at embryonic day (E) 14.5. Furthermore, pituitary specific knockouts of *Jarid2* confirm the hypoplasia of corticotrope and melanotrope lineages and hyperplasia in pituitary stem cells, without affecting the differentiation of *Pou1f1* or *Nr5a1* lineages. With the first-ever pituitary single nucleus (sn) multi-omics at E14.5, we further characterized the RNA expression changes and chromatin landscape reprogramming in pituitary stem cells due to *Jarid2* mutation. Our results suggest that *Jarid2* is a regulator of pituitary stem cell differentiation to POMC lineage in embryonic stages.

## MATERIALS AND METHODS

### Mice

Wild-type CD-1 mice were purchased from Charles River and have been maintained in house. B6;129-*Jarid2^tm1Yskl^*/J (*Jarid2^Fl/Fl^*, RRID:IMSR_JAX:031141) mice were originally generated by Dr. Youngsook Lee (Mysliwiec et al., 2006) by inserting loxP sites flanking exon 3 of *Jarid2*. The strain in our lab was obtained from Jackson Laboratories (RRID:IMSR_JAX:031141, Bar Harbor, ME, United States). Next, germline loss-of-function (LOF) mutation of *Jarid2* (*Jarid2^mut/mut^*) was generated by breeding *Jarid2^Fl/Fl^* to the B6.FVB-Tg (EIIa-cre)C5379Lmgd/J (EIIa-Cre, RRID:IMSR_JAX:003724) mice acquired from Jackson Laboratories. For pituitary-specific LOF mutation, *Jarid2^Fl/Fl^*mice were crossed with *Prop1^T2AiCre^* mice (a kind gift from Dr. Karine Rizzoti (Das et al., 2026), the Francis Crick Institute, United Kingdom) to generate heterozygous *Jarid2^+/Fl^; Prop1^+/Cre^*. The female mice of this genotype were then bred back with *Jarid2^Fl/Fl^*. The *Jarid2^Fl/Fl^; Prop1^+/Cre^*offsprings were designated as *Jarid2* pituitary specific LOF (*Jarid2^PitKO/PitKO^*), and their *Jarid2^Fl/Fl^* littermates were used as controls. Both sexes are included in the analysis.

Mice for wet-lab experiments were maintained at the University of Illinois at Urbana-Champaign animal facility and given food and water *ad libitum*. All animal experimental procedures were approved by the Illinois Institutional Animal Care and Use Committee (IACUC). The *Jarid2^PitKO/PitKO^* mice for single nucleus (sn) multiomics tissue collection were maintained at the University of Michigan according to a protocol approved by Michigan IACUC. The morning of plug identification was designated as embryonic day (E) 0.5, and the day of birth was designated as postnatal day (P) 0.

### Histology and RNAscope *in situ* hybridization

Histology and RNAscope in situ hybridization procedures were performed as previously described (Ge et al., 2024). Briefly, whole embryos at E10.5, E12.5, and E14.5 or whole heads at E16.5 and E18.5 were collected and fixed overnight in 3.7% formaldehyde (Sigma Aldrich) in phosphate-buffered saline (PBS). The samples were then embedded in paraffin and sectioned at 6 μm thickness. For histology, hematoxylin and eosin (H&E) staining was applied to observe pituitary morphology.

For RNAscope *in situ* hybridization, all laboratory glassware was baked to remove RNase contamination. After sectioning, slides were processed with RNAscope Multiplex Fluorescent Reagent Kit v2 (Advanced Cell Diagnostics, ACD) following the manufacturer’s instructions. Probes specific for *Jarid2* were purchased from ACD.

### Immunohistochemistry and cell counting

For immunohistochemistry (IHC) of the nuclear proteins and transcription factors (H3K27me3, SOX2, POU1F1, PAX7, and NR5A1, antibody information in Table 1.), the slides were deparaffinized, rehydrated in graded ethanol washes, and boiled in 10 mM citric acid (pH 6) for 10 minutes for antigen retrieval. Then, the slides were incubated with normal donkey serum diluted in IHC block (3% BSA and 0.5% Triton X-100 in PBS) for 1 hour, followed by overnight incubation at 4℃ in a humidity chamber with primary antibodies diluted to optimal concentration in IHC block. Subsequently, donkey-derived anti-mouse and anti-rabbit secondary antibodies were applied for 1 hour at room temperature to amplify the signal. For proteins that required tertiary amplification (POMC, SOX2, NR5A1, and POU1F1), tertiary antibodies conjugated with streptavidin and fluorophores were then applied for 1 hour at room temperature. Detailed information about the antibodies used and dilution ratios can be found in Table 1. For E16.5 samples, TrueBlack lipofuscin (Biotium) was applied after staining to quench the autofluorescence of red blood cells. All slides were then counterstained with 4′,6-diamidino-2-phenylindole (DAPI; 1:1000; Thermo Fisher Scientific) to stain the nuclei and mounted with an antifade medium (Gonigam et al., 2023). All stained slides were imaged with a fluorescence microscope (Leica DM2560), and all images were processed with Adobe Photoshop 2025.

**Table 1.** Antibody information.

| Antigen | Host | Source | RR_ID | Dilution | Secondary | Tertiary |
| --- | --- | --- | --- | --- | --- | --- |
| H3K27me3 | Rabbit | Cell Signaling Technology | AB_2616029 | 1:400 | Rabbit-Biotin | Strep-Cy3 |
| SOX2 | Rabbit | Millipore | AB_2286686 | 1:2000 | Rabbit-Cy3 | -- |
| MKi67 | Mouse | BD Biosciences | AB_393778 | 1:1000 | Mouse-Cy3 | -- |
| POMC | Rabbit | Agilent Technologies | AB_2936486 | 1:100 | Rabbit-Biotin | Strep-Cy3 |
| POU1F1 | Rabbit | Dr Simon Rhodes | AB_2722652 | 1:500 | Rabbit-Biotin | Strep-Cy3 |
| NR5A1 | Mouse | Invitrogen | AB_2532209 | 1:200 | Mouse-Biotin | Strep-Cy3 |
| PAX7 | Mouse | Developmental Studies Hybridoma Bank | AB_528428 | 1:400 | Mouse-Biotin | Strep-Cy2 |
| Secondary antibodies and tertiary conjugates |  |  |  |  |  |  |
| Antibody | RRID |  | Dilution |  | Source |  |
| Rabbit-Biotin | AB_2340594 |  | 1:250 |  | Jackson ImmunoResearch |  |
| Mouse-Biotin | AB_2340787 |  | 1:250 |  | Jackson ImmunoResearch |  |
| Mouse-Cy3 | AB_2340816 |  | 1:250 |  | Jackson ImmunoResearch |  |
| Rabbit-Cy3 | AB_2313568 |  | 1:250 |  | Jackson ImmunoResearch |  |
| Streptavidin-Cy3 | AB_2337244 |  | 1:250 |  | Jackson ImmunoResearch |  |
| Streptavidin-Cy2 | AB_2337246 |  | 1:250 |  | Jackson ImmunoResearch |  |

To count the immuno-positive cells, three sagittal sections of E14.5 whole embryo at a 10-slide interval (∼ 120 μm) were taken from mid-sagittal to mid-lateral to far-lateral sampling through entire pituitary. The stem cell proportion of the mid-lateral section was quantified by the ratio of area of SOX2-postive cells versus the whole pituitary. For E16.5 embryos, 2 coronal sections spaced 5 slides apart (∼ 60 μm) were counted to minimize location effects. After staining and imaging, ImageJ was employed to count the cells and measure the area of the pituitary in pixels, before converting to area in μm^2^ by the constant of 2.73*2.73 pixels per μm^2^ for 20X images and 5.46*5.46 pixels per μm^2^ for 40X images. All biological replicates were represented as the sum of immuno-positive cell numbers normalized to pituitary areas across all sections.

### Sn multiomics library preparation and sequencing

We dissected 6 pituitary glands from *Jarid2^Fl/Fl^* and 6 pituitary glands from *Jarid2^PitKO/PitKO^* mice at E14.5. Individual Rathke’s pouch samples were flash frozen and stored at −80. Single-cell processing, library prep and next-generation sequencing was carried out in the Advanced Genomics Core at the University of Michigan. Single nuclei suspensions were isolated from flash frozen tissue and subjected to counting on the LunaFx7 automated cell counter (Logos Biosystems) using AO/PI staining before being diluted to 3000-8000 nuclei/µl. Single nuclei libraries were generated using the 10X Genomics Chromium Controller following the manufacturer’s protocol for Multiome (ATAC+GEX) analysis. In brief, transposase was added to the nuclei suspension and incubated, before adding barcoding and RT master mix and loading into the Chromium Controller chip with appropriate gel beads. Following generation of single cell gel bead-in-emulsions (GEMs), reverse transcription of the RNA and barcoding of the fragmented DNA was performed. The resulting product was cleaned and both the cDNA and barcoded, fragmented DNA was amplified. One fraction of this product was used to make the ATAC library via indexing PCR. The other fraction was amplified again to further isolate the cDNA, which then was quantified by Qubit (Invitrogen) and assessed for size on the TapeStation 4200 (Agilent). It then goes through enzymatic fragmentation and a size selection, adapter ligation and indexing PCR. Final library quality was assessed using the LabChip GX Touch HT (Revvity) and libraries were quantified by Qubit. Pooled libraries were then subjected to paired-end sequencing according to the manufacturer’s protocol (Illumina NovaSeq XPlus). BclConvert software (Illumina) was used to generate de-multiplexed Fastq files and the CellRanger Pipeline (10X Genomics) was used to align reads to the mm39 genome and generate count matrices.

### Sn multiomics data processing

For snRNAseq, SoupX (Young and Behjati, 2020) was first employed separately on the control and *Jarid2^PitKO/PitKO^*datasets to estimate and remove the ambient RNA contamination. An additional 10% was added to the estimated contamination fraction to ensure the removal of 95%-98% of contamination. Subsequently, the cleaned data were imported into Seurat 5.2.1 (Hao et al., 2024) to perform all downstream analyses following vignettes provided by the developers. Cells that met these criteria were kept for quality control: nFeature_RNA < 7500, nFeature_RNA > 600, and nCount_RNA < 20000. To further exclude low-quality cells, we applied a data-driven threshold for mitochondrial gene percentage, removing any cells with a percentage exceeding three standard deviations above the mean. Two samples were integrated with the IntegrateLayers() function using canonical correlation analysis (CCA). Normalization, feature selection, scaling, and dimensionality reduction were then performed with default parameters. Next, cell types were annotated based on the expression of previously known markers (Brinkmeier et al., 2025). For differential expressed gene (DEG) analysis, the Wilcoxon test was applied using FindMarkers() with the parameters of logfc.threshold = 0.3 and min.pct = 0.1. Top 50 genes contained in ambient RNA identified by SoupX were removed from DEG analysis. Genes identified with adjusted p-values less than 0.05 were kept as DEGs. For gene ontology (GO) term analysis, the Database for Annotation, Visualization, and Integrated Discovery (DAVID) (Sherman et al., 2024) was used, keeping only GO terms with p-values < 0.05. Pseudotime trajectory was constructed by slingshot package (Street et al., 2018). Mean pseudotime comparison between genotypes was conducted by permutation test and comparison on the distribution of pseudotime was conducted by PrgressionTest (Global = TRUE) in condiments package (De Bézieux et al., 2021).

MAGIC analysis was utilized to infer TF enrichment of the DEGs of snRNAseq based on the ENCODE archive Chromatin Immunoprecipitation Sequencing (ChIP-seq) (Roopra, 2020). Analysis was conducted using _magic_3.py according to Dr. Roopra’s instructions, keeping general transcription factors and RNA polymerase in the analysis and zero ChIP values (-g y -z y).

For snATACseq, Signac 1.14.0 (Stuart et al., 2021) was utilized for all analyses following vignettes provided by the developers. Before merging control and *Jarid2^PitKO/PitKO^* samples, a common peak set was first generated, including peaks with widths between 20 and 10,000 bp. Peaks located outside standard chromosomes were also excluded. For quality control, the following standards were applied: nCount_ATAC > 500, nCount_ATAC < 100,000, pct_reads_in_peaks > 15, blacklist_ratio < 0.05, and TSS.enrichment > 3. Normalization and linear dimensionality reduction were subsequently conducted using parameters from Signac’s vignette “merging objects”. Next, the cell types were annotated by transferring the annotations from the snRNAseq with FindTransferAnchors() and TransferData(). Gene activity was calculated by the GeneActivity() function and normalized using ‘LogNormalize’ method and scaled to the median nCount of gene activity. To identify the genes that had overall gene activity change, the Model-based Analysis of Single-cell Transcriptomics (MAST) model (Finak et al., 2015) was used with latent variables set to cell types and nCount_ATAC. To analyze TF binding motif activity, motif information was first added from the JASPAR 2020 database (Fornes et al., 2020). Motif activity per cell was then calculated with chromVAR (Schep et al., 2017). For DEG analysis of motif activity, the logistic regression (LR) framework was used as recommended by the developers with parameters of logfc.threshold = 0.1, min.pct = 0.1, latent.vars = “nCount_ATAC”, and mean.fxn = rowMeans.

### Postnatal RNA isolation and quantitative real-time polymerase chain reaction (qRT-PCR)

Total RNA of P0 mouse pituitaries was extracted with the RNAqueous Total Micro Kit (Thermo Fisher Scientific) according to the manufacturer’s protocol. Extracted RNA was reverse transcribed to complementary DNA (cDNA) using the Protoscript First Strand cDNA Synthesis Kit (New England Biolabs). For qRT-PCR, all cDNA samples were amplified with platinum Taq DNA polymerase (Fisher Scientific) in a SYBR green (Fisher Scientific) containing in-house mastermix (Weis et al., 2023) using primers specific for *Meis2* (Forward: 5’-TTCCAGCATCTCACACACCC-3’, Reverse: 5’-TCACTGCTCGATTTGACTGGT-3’) and the housekeeping gene *Gapdh*. *Meis2* expression level was normalized to *Gapdh* using the 2^-ΔΔCt^ method (Nantie et al., 2014).

### Statistical analysis

All data were represented as mean ± standard error of mean (SEM). Three or more biological replicates were included for each experiment. For experiments comparing two groups, Student’s t-test was used to determine statistical significance. E16.5 data with both sex and genotype variables were first analyzed with one-way analysis of variance (ANOVA) to determine whether there was a sex difference, followed by multiple comparison. No sex differences were discovered in this study, so all data were combined between sexes. P-values less than 0.05 were considered statistically significant. To reduce false positives in sn multiomics, more stringent thresholds were employed with single-gene expression comparisons using Wilcoxon’s rank sum test. Genes with false discovery rate adjusted p-values < 0.001 were considered statistically significant. All analyses and calculations were performed with GraphPad Prism 8.2.1 or R 4.4.2.

### Use of AI

ChatGPT 5.4, Claude 4.7, and Illinois flagship chatbot were used during the preparation of this manuscript. The usage of AI is limited only to the generation of code and simple language editing to improve accuracy and clarity.

## RESULTS

### *Jarid2* is present ubiquitously while H3K27me3 has a distinct distribution pattern

To determine the localization of *Jarid2* mRNA, we performed RNAscope in-situ hybridization on mouse embryos at critical timepoints of pituitary development. *Jarid2* mRNA is ubiquitously present in the pituitary from E10.5, E12.5, E14.5 to E18.5 (Fig. 1A-D), with no distinct enrichment in PL, IL, or AL. To further characterize the distribution pattern of PRC2 subunits, we analyzed our control snRNAseq dataset at E14.5 for the mRNA distribution of the core PRC2 subunits (*Ezh1/2*, *Eed*, *Suz12*, and *Rbbp4/7*) and facultative subunits (*Jarid2*, *Aebp2*, *Mtf2*, *Phf1*, *Phf19*, *Epop*, *Lcor*, and *Lcorl*; Fig S1G-N). All the PRC2 core components are broadly expressed in mouse pituitary at E14.5 (Fig. S1A-F). *Jarid2* mRNA is present in all pituitary and hypothalamus clusters identified in snRNAseq (Fig. S1G), alongside the other PRC2.2 subunit *Aebp* (Fig. S1H). In the PRC2.1 subcomplex, while *Mtf2*, *Lcor* (encoding PALI1) and *Lcorl* (encoding PALI2) are ubiquitously expressed (Fig. S1L, Fig. S1M-N), *Phf1* (Fig. S1J) and *Epop* (Fig. S1L) are not highly expressed in any of the clusters. Interestingly, *Phf19* is enriched only in the rostral tip (RT) thyrotropes (Fig. S1K), the first hormone-secreting cell type to differentiate in embryonic pituitary.

**Fig. 1.**
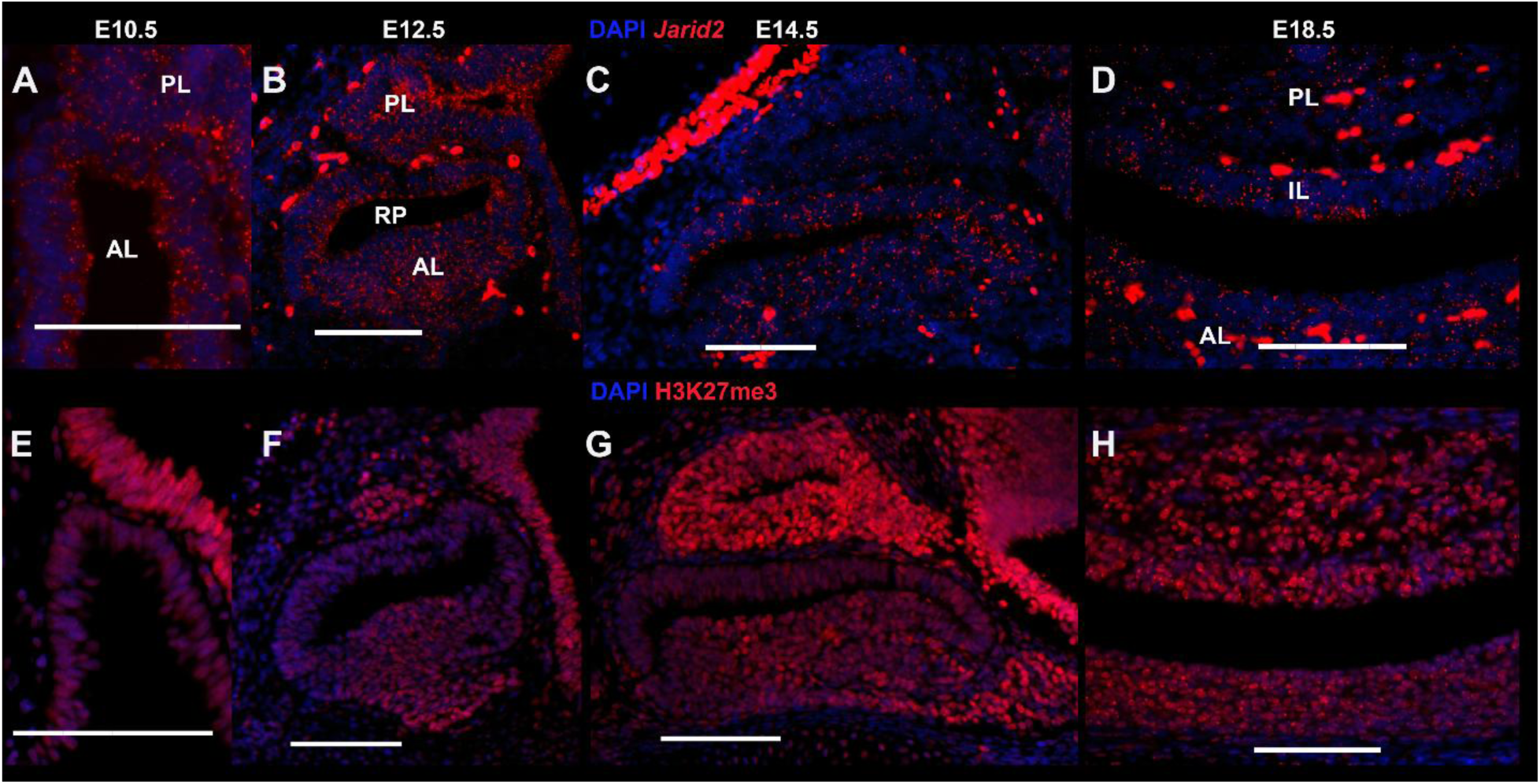
Localization of *Jarid2* mRNA and H3K27me3 in embryonic pituitary from E10.5 to E18.5. (A-D) RNAscope *in situ* hybridization shows *Jarid2* mRNA is ubiquitously expressed at embryonic day (E) 10.5(A), E12.5(B), E14.5(C), and E18.5(D). (E-H) Trimethylation at lysine 27 of histone H3 (H3K27me3) is high in posterior lobe and low in anterior lobe at E10.5(E). The level of H3K27me3 is increased in differentiated anterior lobe at E12.5(F) and E14.5(G). H3K27me3 is ubiquitous at E18.5(H). AL: anterior lobe; IL: intermediate lobe; PL: posterior lobe; RP: Rathke’s pouch. Scale bar: 100 μm.

Based on the ubiquitous pattern of *Jarid2* and PRC2 core components, we hypothesized that H3K27me3 is also ubiquitously distributed throughout embryonic pituitary development. To test that, we conducted immunohistochemistry (IHC) of H3K27me3 at the previously mentioned developmental timepoints. To our great surprise, H3K27me3 has a distinct distribution pattern: at E10.5, H3K27me3 level is relatively high in the forming PL compared to the rudimentary Rathke’s pouch (Fig. 1E). Subsequently at E12.5 and E14.5, H3K27me3 remains high in PL (Fig. 1F,G). Notably, the stem cells residing around the cleft have low H3K27me3 levels and the differentiated RT has elevated H3K27me3. At E18.5, the difference in H3K27me3 between AL and PL disappears and H3K27me3 becomes widespread in the developing pituitary. Our result suggests that H3K27me3 bulk levels increase as pituitary stem cells differentiate independently of the expression of PRC2 complex subunits, suggesting that H3K27me3 may contribute to pituitary stem cell differentiation.

### Germline loss of *Jarid2* leads to hyperplasia of pituitary stem cells

To uncover the role of *Jarid2* in pituitary development, we generated a germline loss-of-function (LOF) mutation mouse (*Jarid2^mut/mut^*) by crossing *Jarid2* floxed mice with EIIa-cre mice. First, hematoxylin and eosin staining with the mid-sagittal section of *Jarid2^mut/mut^*and their wildtype littermates at E14.5 did not show any observable difference (Fig. 2A,D). To capture the morphology of the pituitary along the left-right axis, we stained three sections every 120 μm from the same embryo ranging from mid-sagittal, mid-lateral, to far-lateral (Fig. 2A-F). *Jarid2^mut/mut^* mid-lateral sections have a hyperplastic Rathke’s pouch (Fig. 2B,E). To validate the expanded tissues have stem cell identity, we performed IHC on the mid-lateral slides with SOX2 and MKi67 antibodies (Fig. 2G,H). We found that the SOX2+ stem cell population expands in *Jarid2^mut/mut^*mid-lateral sections, with no observable change in their proportion of proliferation. Further, quantification of the proportion of stem cell area confirmed that there is a significant increase in cleft stem cells in *Jarid2^mut/mut^*mid-lateral pituitary at E14.5 (Fig. 2I). These results suggest that *Jarid2* is important for limiting the number of pituitary stem cells at E14.5 and the effect is restricted to the mid-lateral plane.

**Fig. 2.**
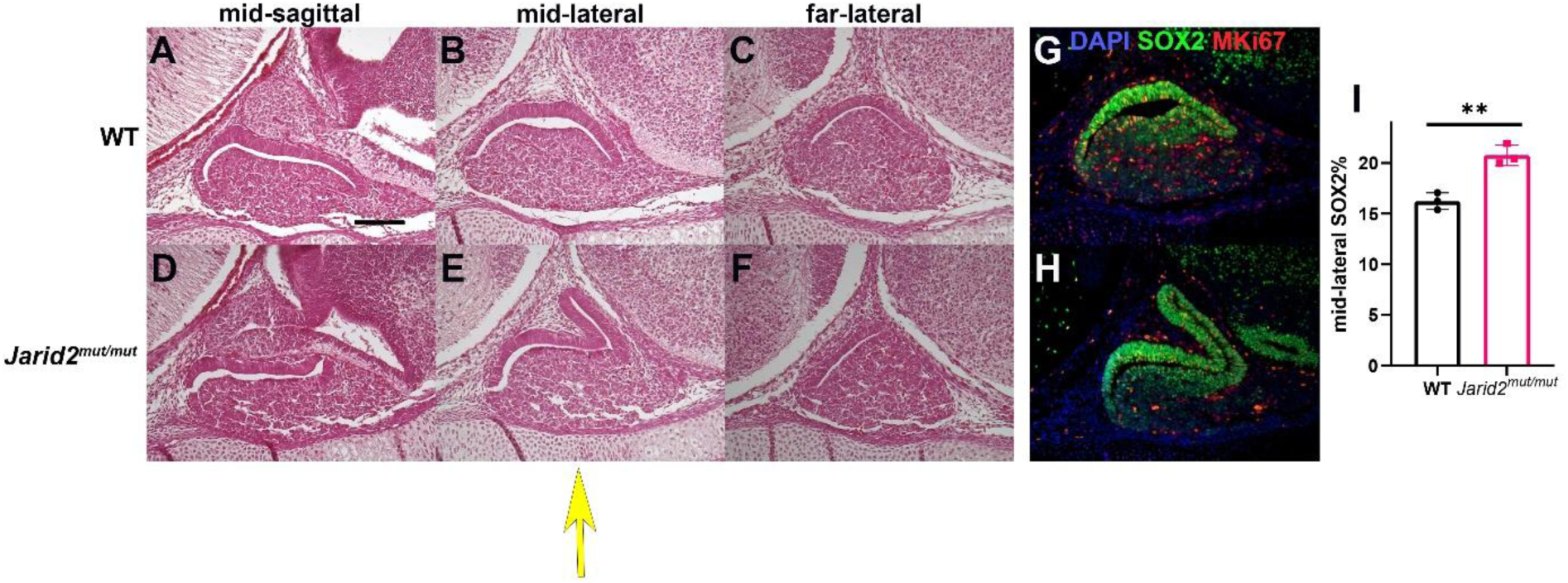
Germline loss of *Jarid2* leads to mid-lateral pituitary stem cell hyperplasia at E14.5. (A-F) Hematoxylin and Eosin staining on mid-sagittal, mid-lateral, and far-lateral sections of wildtype(A-C) and *Jarid2^mut/mut^*(D-F) at E14.5. Yellow arrow signifying change in mid-lateral section. (G,H) Immunohistochemistry shows the level of SOX2 (Green) and MKi67 (Red) in mid-lateral section of wildtype(G, WT) and *Jarid2^mut/mut^*(H). (I) Quantification shows that the proportion of stem cell area in pituitary is significantly increased, N = 3, “**” indicates that p-value < 0.01. Scale bar: 100 μm.

### Germline loss of *Jarid2* causes corticotrope hypoplasia without affecting POU1F1 progenitors

Given the stem cell hyperplasia in *Jarid2^mut/mut^* at E14.5, we then investigated the function of *Jarid2* in stem cell differentiation by IHC of lineage markers POMC and POU1F1 on *Jarid2^mut/mut^*e14.5 embryos. There is a strong reduction in the number of POMC+ cells in *Jarid2^mut/mut^*(Fig. 3A,B), both in AL and IL. Quantification on the anterior lobe side confirms a significant reduction in corticotrope cell number (Fig. 3G).This result suggests that *Jarid2* may regulate the differentiation of both corticotrope and melanotrope cells. Nonetheless, POU1F1 staining shows that POU1F1+ cell number is not observably changed between *Jarid2^mut/mut^*and wildtype (Fig. 3C,D), suggesting a potential POMC-lineage specific role of *Jarid2* at E14.5.

**Fig. 3.**
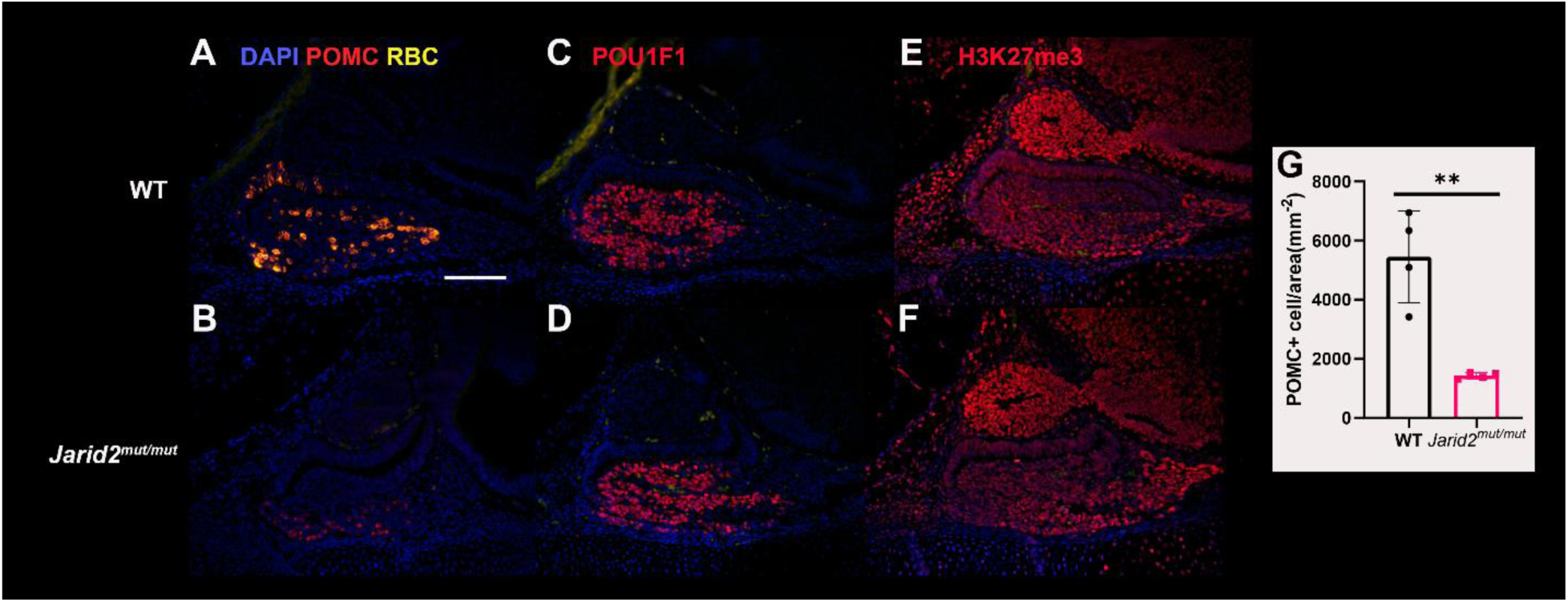
Germline loss of *Jarid2* affects *Pomc* lineage formation without affecting *Pou1f1* lineage or H3K27me3 bulk levels at E14.5. (A, B) Immunohistochemistry shows that POMC+ cells are significantly reduced in *Jarid2^mut/mut^* (B) compared to its wildtype littermate(A). (C, D) POU1F1+ cell number is similar between wildtype (C) and *Jarid2^mut/mut^* (D). (E-F) Localization of H3K27me3 measured by immunohistochemistry is not changed in wildtype(E) compared to *Jarid2^mut/mut^* (F). (G) Quantification of POMC+ cell number normalized to anterior pituitary area, N = 4, “**” indicates that p-value < 0.01. Scale bar: 100 μm.

As shown in Fig. 1G, H3K27me3 has an increasing gradient as pituitary stem cells differentiate. Therefore, we hypothesize that *Jarid2* is regulating pituitary stem cell differentiation by modulating the bulk level of H3K27me3. However, IHC on H3K27me3 did not indicate detectable changes in the bulk level of H3K27me3 (Fig. 3E,F). This result is consistent with previous experiments in mouse embryonic stem cells that *Jarid2* mutations do not cause a global de-repression of PRC2 or measurable change in H3K27me3 level (Landeira et al., 2010). To summarize, we identified mid-lateral stem cell hyperplasia and overall corticotrope hypoplasia in *Jarid2^mut/mut^* at E14.5, with no effect on POU1F1 cell number nor H3K27me3 bulk level.

### Pituitary-specific loss of *Jarid2* selectively inhibits the POMC lineage

Since *Jarid2^mut/mut^* is lethal around E15.5 (Takeuchi et al., 1995). it is impossible to observe the differentiation of all pituitary lineages, especially the *Nr5a1* lineage with this model. To further characterize pituitary lineage differentiation and determine pituitary intrinsic effects, we crossed *Jarid2^Fl/Fl^* with *Prop1*-Cre mice kindly provided by Dr. Karine Rizzoti (Das et al., 2026), generating pituitary-specific LOF mutation of *Jarid2* (*Jarid2^PitKO/PitKO^*). Single nucleus RNAseq on E14.5 embryos of *Jarid2^PitKO/PitKO^*and their *Jarid2^Fl/Fl^* littermates validated deletion of exon 3 of *Jarid2* in pituitary cells of *Jarid2^PitKO/PitKO^*(Fig. S2). Interestingly, there seems to be an upregulation of the overall *Jarid2* mRNA. This result indicates that although the deletion of *Jarid2* exon 3 is predicted to generate a frameshift mutation and a truncated non-functional 15-amino-acid JARID2 peptide (Mysliwiec et al., 2006), this *Jarid2* mRNA is not subjected to nonsense mediated decay (Lykke-Andersen and Jensen, 2015).

Using this model, we first examined corticotrope differentiation at E14.5. There is a subtler but significant reduction of POMC+ cell number in the anterior lobe (Fig. 4A,B,E). This result reveals that *Jarid2* regulates corticotrope differentiation in a pituitary-intrinsic manner. Consistent with *Jarid2^mut/mut^*, POU1F1+ cell number remains similar between *Jarid2^Fl/Fl^* and *Jarid2^PitKO/PitKO^*(Fig. 4C,D), suggesting *Jarid2* has a selective role in the formation of the *Pomc* lineage rather than the *Pou1f1* lineage.

**Fig. 4.**
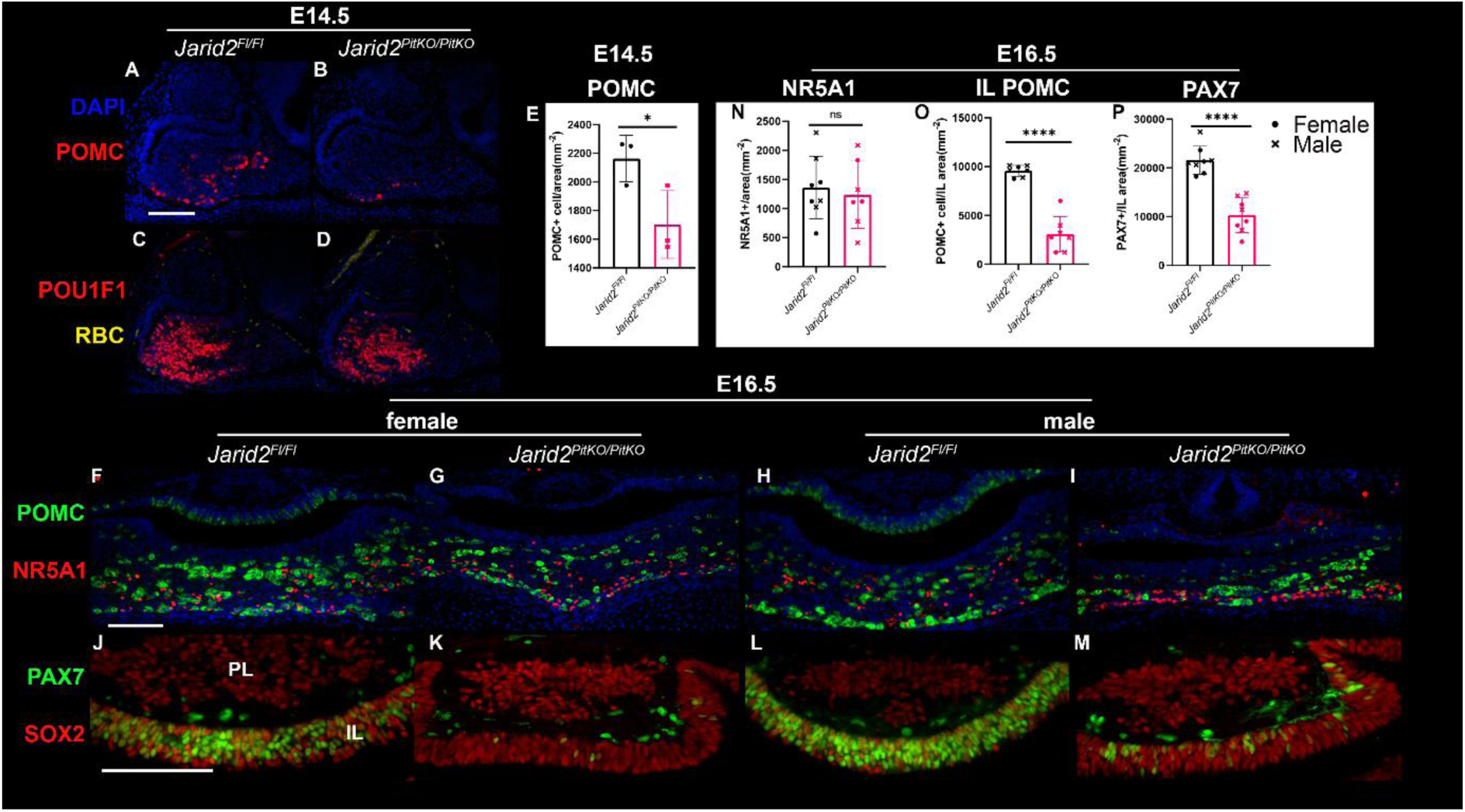
Pituitary specific loss of *Jarid2* leads to reduction in POMC and PAX7 without affecting NR5A1 progenitor numbers at E14.5 and E16.5. (A,B) POMC+(red) cells detected by immunohistochemistry at E14.5 are reduced in *Jarid2^PitKO/PitKO^*(B) compared to *Jarid2^Fl/Fl^* littermate(A). (C, D) POU1F1 staining of *Jarid2^Fl/Fl^* (C) and *Jarid2^PitKO/PitKO^*(D). (E) Quantification of POMC+ cell number per AL area at E14.5, N = 3, “*” indicates p-value < 0.05. (F-I) POMC(green) and NR5A1(red) double staining in both sexes at E16.5 in *Jarid2^Fl/Fl^* (F,H) and *Jarid2^PitKO/PitKO^* (G,I). (J-M) PAX7 (green) and SOX2 (red) double staining in IL in *Jarid2^Fl/Fl^* (J,L) and *Jarid2^PitKO/PitKO^* (K,M) in both sexes showing a reduction in PAX7 cell number. (N) Quantification of NR5A1+ cell per AL area, “ns” means not significant. (O) Reduction of POMC+ cell number in *Jarid2^PitKO/PitKO^* in IL at E16.5. (P) Reduction of PAX7+ cell number at in *Jarid2^PitKO/PitKO^* E16.5. N = 7-8 (mixed sex), “****” indicates P < 0.0001. “●” indicates female. “X” indicates male. Scale bar is 100 μm.

To further reveal the role of *Jarid2* for all pituitary lineages, we investigated pituitary differentiation in *Jarid2^Fl/Fl^* and *Jarid2^PitKO/PitKO^* at E16.5 with IHC. We first captured the onset of the gonadotrope lineage by staining with NR5A1(Fig. 4F-I) and found no significant difference between sexes. In a mixed-sex population, we identified no difference in NR5A1+ gonadotrope progenitor cell number between *Jarid2^Fl/Fl^* and *Jarid2^PitKO/PitKO^*littermates (Fig. 4N). From IHC of POMC at E16.5, we observed that there is a reduction in POMC+ cell number in the IL of *Jarid2^PitKO/PitKO^* E16.5 embryos (Fig. 4F-I). To better understand the variable reduction of melanotrope cell number, we quantified POMC+ cell number normalized to IL area and discovered that the number of POMC+ melanotrope cells is significantly reduced in *Jarid2^PitKO/PitKO^* (Fig. 3O). Together, these results show that *Jarid2* regulates the differentiation of both corticotropes and melanotropes without affecting gonadotrope progenitors or POU1F1+ somatotrope/thyrotrope/lactotrope progenitors.

*Pax7* is a pioneer TF that specifies melanotrope cell fate over corticotropes by regulating chromatin accessibility (Budry et al., 2012; Mayran et al., 2018). To further investigate how *Jarid2* regulates the differentiation of the POMC lineage melanotropes, we co-stained pituitary sections with the stem cell marker SOX2 and PAX7 in *Jarid2^PitKO/PitKO^* embryos. At E16.5, SOX2 is expressed in every cell of the IL and every PAX7+ cell overlaps with SOX2 (Fig. 4J,L), consistent with previous studies (Budry et al., 2012). PAX7 is strongly reduced in the IL of *Jarid2^PitKO/PitKO^* at E16.5 (Fig. 4K,M), further confirmed by a statistically significant quantification (Fig. 4P). Unlike *Tbx19^-/-^*pituitaries where *Pax7* is limited to the ventral/luminal side (Budry et al., 2012), *Jarid2^PitKO/PitKO^* pituitaries exhibit a global reduction of PAX7+ cells across the IL (Fig. 4J-N). It is worth mentioning that *Jarid2* LOF mutation does not completely deplete POMC or PAX7 in the IL, suggesting that *Jarid2* is not the sole determinant of POMC lineage commitment. Collectively, these results indicate that *Jarid2* specifically regulates the differentiation of POMC lineage during embryonic pituitary development.

### Pituitary specific loss of *Jarid2* induces pituitary stem cell hyperplasia

Since *Jarid2^PitKO/PitKO^* showed a defect in POMC lineage differentiation, we hypothesized that *Jarid2* is regulating the lineage commitment of pituitary stem cells. Therefore, we characterized the number of pituitary stem cells in the AL by IHC with SOX2 antibodies. At first examination, *Jarid2^PitKO/PitKO^*pituitaries have a spectrum of stem cell hyperplasia in both sexes (Fig. 5A-F). Some sections look similar to their *Jarid2^Fl/Fl^* littermates (Fig. 5B,E), while others have a severe hyperplastic and dysmorphic cleft (Fig. 5C,F). In severely affected individuals (Fig. 5C), the mid-sagittal plane may seem normal compared to *Jarid2^Fl/Fl^*, whereas the “mid-lateral” plane has an expanded cleft, potentially explaining the mid-lateral specific hyperplasia we observed in *Jarid2^mut/mut^* E14.5 (Fig. 2G,H). To confirm stem cell hyperplasia despite the variability, we quantified the number of pituitary stem cells residing at AL cleft or parenchyma in two sections with a 60-μm-interval in each individual. Results showed that there is a significant increase in the total number of SOX2+ stem cells (Fig. 5I), involving increase in both cleft stem cells (Fig. 5G) and parenchyma stem cells (Fig. 5H). In summary, these results suggest that *Jarid2^PitKO/PitKO^* causes retention of pituitary stem cells due to inability to differentiate into the POMC lineage at E16.5.

**Fig. 5.**
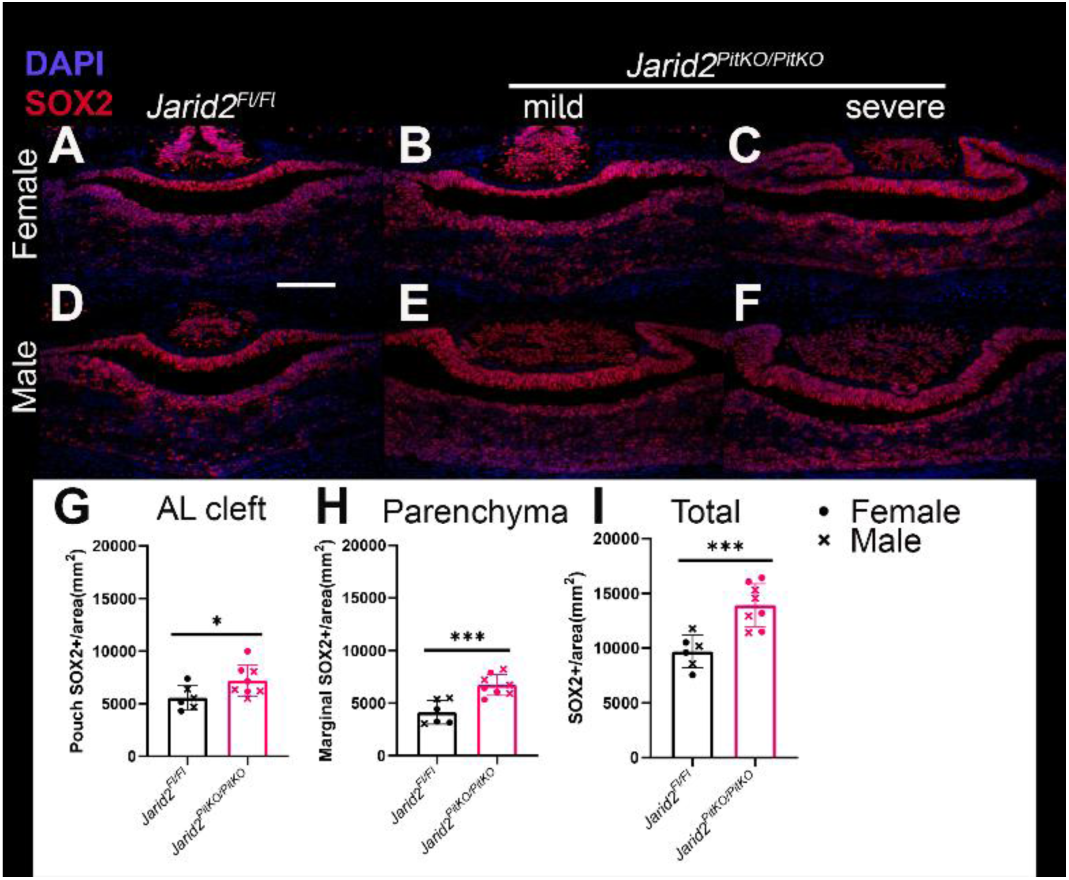
Pituitary specific loss of *Jarid2* causes pituitary stem cell hyperplasia at E16.5. (A-F) Immunohistochemistry probing pituitary stem cell marker SOX2 (red) shows variable severity of stem cell hyperplasia in *Jarid2^PitKO/PitKO^* (B,C,E,F) in both sexes compared to *Jarid2^Fl/Fl^* (A,D). (G-I) Quantification of stem cell number demonstrates an increase in anterior side cleft (G), pituitary parenchyma (H), and total number (I). N = 6-8. “*” indicates P < 0.05. “***” indicates P < 0.0001. “●” indicates female. “X” indicates male. Scale bar is 100 μm.

### SnRNAseq reveals alterations in gene expression profile in pituitary progenitors due to loss of *Jarid2*

To begin to determine the mechanism of how *Jarid2^PitKO/PitKO^* impacts pituitary stem cells and POMC lineage differentiation, changes in RNA expression and chromatin accessibility at the single-nucleus level were assessed by sn-multiomics, snRNAseq and snATACseq on nuclei isolated from mixed-sex E14.5 pituitaries from *Jarid2^PitKO/PitKO^*or *Jarid2^Fl/Fl^* littermates. Initial dimensionality reduction shows that *Jarid2^PitKO/PitKO^* and *Jarid2^Fl/Fl^*have similar UMAP visualizations with no evident batch effect (Fig. 6A). We then annotated cell identities (Fig. 6B) with previously established cell type markers in E14.5 pituitaries (Brinkmeier et al., 2025). We identified proliferating progenitors (PP), *Sox2*+ progenitors (*Sox2* Prog), *Prop1*+ progenitors (*Prop1* Prog), *Prop1/Pou1f1* double positive progenitors (*Prop1-Pou1f1*), *Pou1f1* progenitors (*Pou1f1* Prog), *Pou1f1* lineage committed cells (*Pou1f1* Com, marked by *Pou1f1* and *Isl1*), corticotrope progenitors (Cort), thyrotrope progenitors (Thyro), and rostral tip thyrotropes (RT Thyro). Results suggest that *Jarid2^PitKO/PitKO^* and *Jarid2^Fl/Fl^* have overall similar cell type compositions (Fig. 6C).

**Fig. 6.**
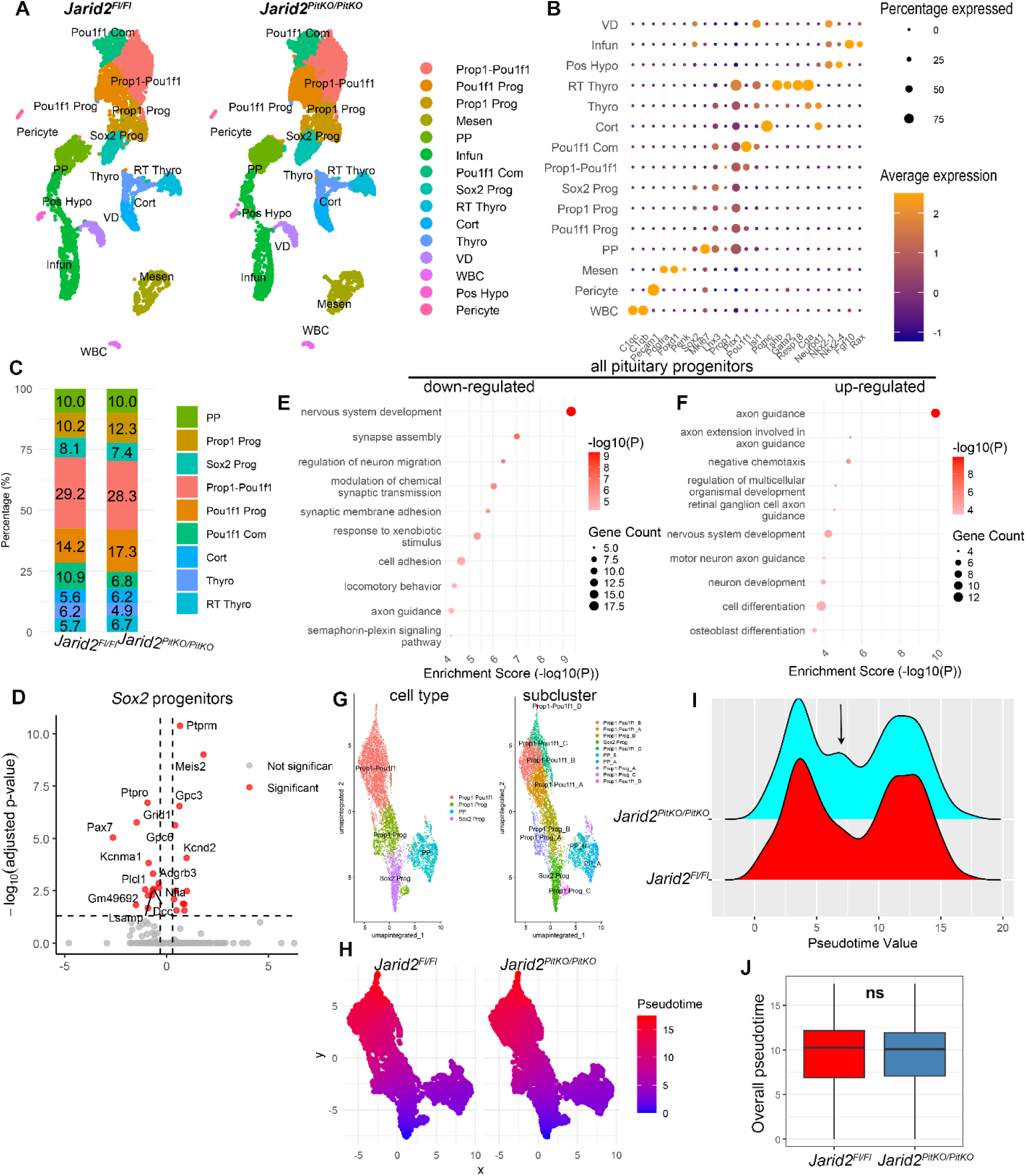
Pituitary specific loss of *Jarid2* alter the single nucleus RNAseq (snRNAseq) expression profile of pituitary progenitors at E14.5. (A) UMAP plot showing similar cluster distribution between *Jarid2^Fl/Fl^* and *Jarid2^PitKO/PitKO^*. Cort: corticotrope; Infun: infundibulum; Mesen: mesenchyme; Pos Hypo: posterior hypothalamus; *Pou1f1* Com: *Pou1f1* lineage committed cells; *Pou1f1* Prog: *Pou1f1* progenitors; PP: proliferating progenitors; *Prop1-Pou1f1*: *Prop1/Pou1f1* double positive progenitors; *Prop1* Prog: *Prop1* progenitors; RT Thyro: rostral tip thyrotrope; *Sox2* Prog: *Sox2* progenitors; Thyro: thyrotrope; VD: ventral diencephalon; WBC: white blood cells. (B) Dot plot showing the expression of marker genes in the clusters. (C) Stacked box plot showing *Jarid2^PitKO/PitKO^* exhibits overall similar clustering with control. (D) Volcano plot of differentially expressed genes (DEG) in *Sox2* Prog. Red dots indicate significant DEGs and grey dots indicate genes that are not significant. (E,F) Dot plot showing gene ontology (GO) term enrichment for down-regulated (E) and up-regulated (F) DEGs in all pituitary progenitors (PP, *Sox2* Prog, *Prop1* Prog, and *Prop1-Pou1f1*). (G) UMAP plot showing the sub-clustering of pituitary progenitors. (H,I) Pseudotime analysis presented in UMAP (H) or ridge plot (I), arrow indicating a bulge in pseudotime distribution.(J) Box plot showing a similar average pseudotime between *Jarid2^Fl/Fl^* and *Jarid2^PitKO/PitKO^*.

We then performed differentially expressed gene (DEG) analysis to capture RNA expression alterations at the cluster level. Drawing from the increase of SOX2+ cells in embryos (Fig. 2,5), we first focused on the *Sox2* Prog cluster (Fig. 6D). Up-regulated genes include *Gpc3*, a surface protein whose mutation leads to pituitary hormone anomaly in patients (Bu et al., 2021), and *Meis2*, a TF whose LOF is reported to cause cleft palate in patients (Zhang et al., 2021) by regulating the development of neural crest (Machon et al., 2015). Interestingly, we observed a significant downregulation of *Pax7* RNA in *Jarid2^PitKO/PitKO^ Sox2* Progenitors (P = 2.7*10^-10^), consistent with a reduction in PAX7 immuno-positive cell number in *Jarid2^PitKO/PitKO^* at E16.5 (Fig. 4P).

Since *Sox2* and *Prop1* are both well-established pituitary stem cell markers (Pérez Millán et al., 2024), we focused on *Sox2* or *Prop1* containing pituitary progenitors (*Sox2* Prog, *Prop1* Prog, PP, and *Prop1-Pou1f1*) to fully capture changes in all pituitary progenitors. Gene ontology (GO) term analysis of down-regulated genes in pituitary progenitors uncovered enrichment in genes related to nervous system development, neuron migration, and cell adhesion, including *Reln*, *Ncam1*, *Nrg1,* and *Syt7* (Fig. 6E). There is also an enrichment in the semaphoring-plexin signaling pathway (including *Sema6d*, *Plxna2*, *Plxnc1*, and *Plxna4*). For upregulated genes in *Jarid2^PitKO/PitKO^* pituitary progenitors, there is an enrichment for axon guidance related genes, which includes *Mef2c*, *Sema3d*, *Sema3a*, *Slit2*, *Slit3*, and *Nrcam* (Fig. 6F). It has been suggested recently that these signaling molecules may play an important role in pituitary development (Kövér et al., 2026). Notably, a few genes related to modulation of the Wnt signaling pathway are also altered, with *Tcf7l2* and *Lgr4* increasing (Table S2) and *Dkk2* decreasing (Table S1). It is worth noting that few changes are identified in the corticotrope cluster (Fig. S4G), suggesting that corticotrope progenitors may not be the primary target of JARID2.

To further understand the role of *Jarid2* in regulating the developmental dynamics of pituitary progenitors, we extracted and subclustered the pituitary progenitors (Fig. 6G). Next, we conducted pseudotime analysis with the slingshot package (Street et al., 2018) and established the pseudotime progression of pituitary progenitors (Fig. 6H-J). Interestingly, there appears to be one additional peak in *Jarid2^PitKO/PitKO^* between the two peaks identified in *Jarid2^Fl/Fl^* (Fig. 6I), indicating a potential arrest in developmental trajectory. Although we discovered no significant shift in overall average pseudotime (Fig. 6J), there is a significant change in pseudotime progression, further confirming a shift in pituitary progenitor developmental dynamics induced by *Jarid2^PitKO/PitKO^*. Taken together, snRNAseq suggests that *Jarid2^PitKO/PitKO^* alters the expression profile of pituitary stem cells, represses *Pax7* expression, and shifts the developmental dynamics of pituitary progenitors.

### *Jarid2* reprograms the chromatin accessibility landscapes in pituitary stem cells

Using snATACseq, we determined changes in the chromatin accessibility landscapes induced by *Jarid2^PitKO/PitKO^* in E14.5 pituitaries. We first annotated the snATACseq dataset with cell identities established in the snRNAseq (Fig. 7A,B). Interestingly, PP do not have a distinct snATACseq cluster, which may indicate that the proliferation program occurs at the level of transcription rather than chromatin accessibility level (Fig. 7B). For other pituitary progenitors, *Prop1-Pou1f1* cluster has two separate ATAC clusters and there is one mixed-identity pituitary stem cell cluster (labeled PSC in the UMAP). The annotation transferred from snRNAseq is confirmed by enrichment of previously known markers (Fig. S3). To obtain a broad understanding of the overall chromatin accessibility, we averaged all the signals within the gene body and promoter region with gene activity algorithm in Signac (Stuart et al., 2021). Using the MAST algorithm with latent variable set to cell type (Finak et al., 2015), we uncovered genes that have changed accessibility independently of cell types in embryonic pituitary (Fig. 7C). Results show that *Jarid2* mutation is primarily associated with increased chromatin accessibility within the pituitary, indicating that *Jarid2* is mainly serving as an epigenetic repressor in embryonic pituitary. To better understand how changes in chromatin accessibility correlate with TF binding activity, we utilized chromVAR algorithm to calculate the gain or loss in chromatin accessibility in the peaks containing known TF binding motifs (Schep et al., 2017). We primarily focused on *Sox2* progenitors and observed an upregulation of TEAD family (TEAD1/2/3/4) and SIX family TF binding motifs (SIX1/2; Fig. 7D,F). TEAD family TFs are downstream effectors of Hippo signaling pathway, whose activation is essential for SOX2+ pituitary stem cell activity (Lodge et al., 2019). Particularly, *Tead2* is shown to be expressed strongly in developing pituitary (Lodge et al., 2016) and is upregulated in neonatal and tumorigenic pituitary stem cells (Cox et al., 2017). We observed an upregulation of TEAD2 binding motif in *Sox2* Prog (Fig. 7F), potentially contributing to the stem cell hyperplasia in the pituitary. SIXs and TEADs are also reported to be enriched in pituitary stem cells in the consensus pituitary atlas (Kövér et al., 2026). In other pituitary progenitor clusters, *Prop1* progenitors also have an elevated level of SIX TF binding motif and *Prop1-Pou1f1*_1 cluster have elevated level of TCF7L2 and LEF1 (Table S6).

**Fig. 7.**
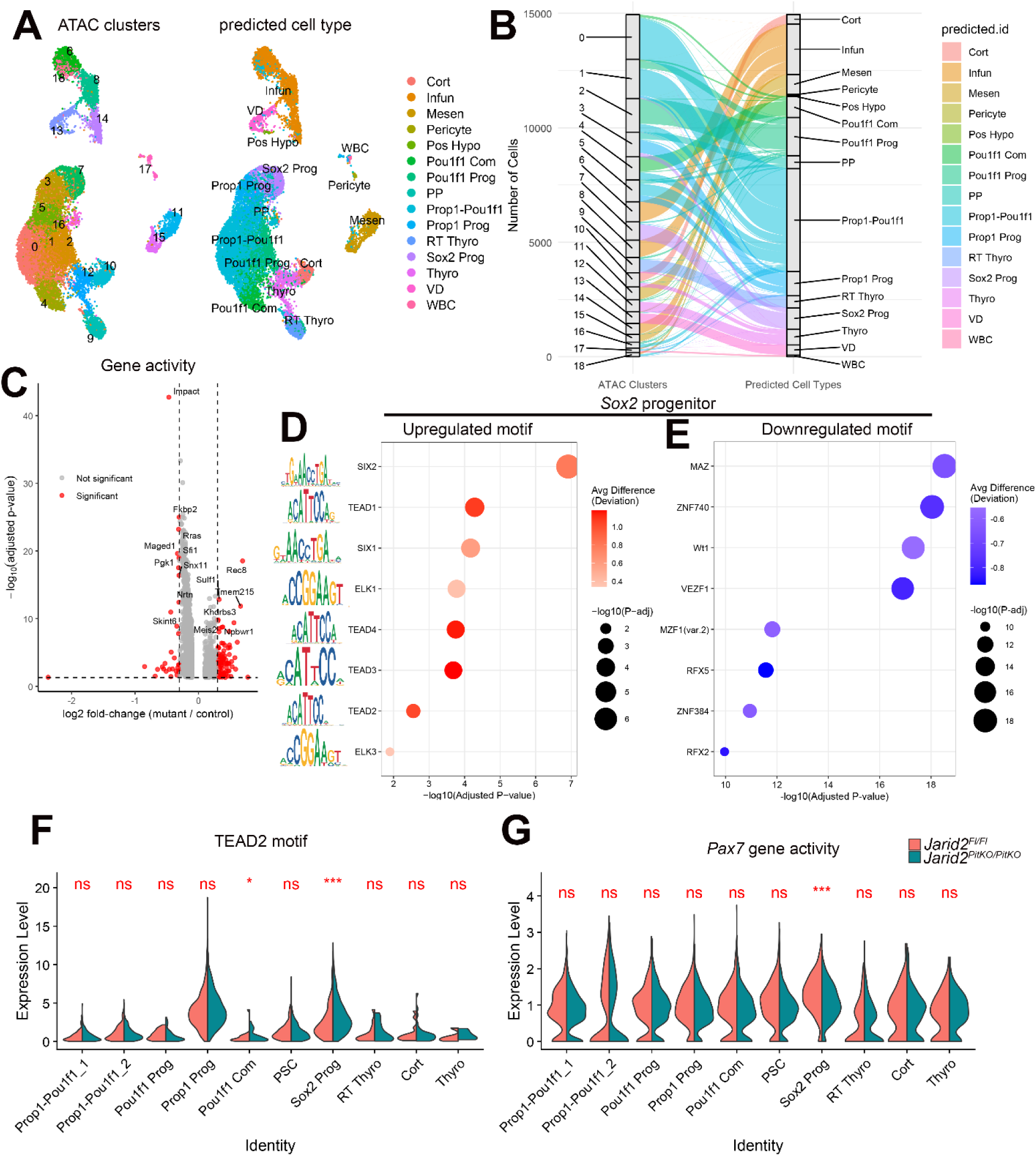
*Jarid2* remodels chromatin accessibility in *Sox2* progenitors at E14.5. (A,B) Cluster annotation of single nucleus Assay for Transposase-Accessible Chromatin with sequencing (snATACseq) predicted from snRNAseq in UMAP (A) and sankey (B) plot. (C) Volcano plot showing the overall difference between *Jarid2^Fl/Fl^* and *Jarid2^PitKO/PitKO^* of snATACseq aggregated chromatin accessibility gene activity identified by MAST. Red dots indicate significant genes, and grey dots indicate non-significant genes. (D,E) Motif enrichment analysis of transcription-factor-associated up-regulated (D) or down-regulated(E) chromatin accessibility in *Sox2* Prog identified by chromVAR. (F) Violin plot of the TEAD2 motif change between *Jarid2^Fl/Fl^* and *Jarid2^PitKO/PitKO^* in all pituitary clusters. (G) Violin plot of *Pax7* gene activity. “ns” indicates not significant. “*” indicates false discovery rate adjusted p-value (Padj) < 0.001. “***” indicates Padj < 1*10^-5^.

The TF binding motif for RFX2 is down-regulated in *Jarid2^PitKO/PitKO^ Sox2* Prog (Fig. 7E). These TFs are related to pituitary adenomas but have unknown function in pituitary development (Raimondi et al., 2019; Wang et al., 2022)Two TFs that are essential for pituitary differentiation (Edwards and Raetzman, 2018), LHX3 and PITX1, are downregulated in multiple clusters of *Jarid2^PitKO/PitKO^* cells (LHX3 in *Prop1* Prog, *Sox2* Prog, *Pou1f1* Prog, and *Prop1-Pou1f1*_1/2; PITX1 in *Prop1* Prog, *Pou1f1* Prog, PSC, and *Prop1-Pou1f1*_2) (Table S5). The binding motif of INSM1, an Zn finger TF essential for pituitary differentiation (Welcker et al., 2013), is downregulated in all pituitary clusters (Table S5). Taken together, these results suggest that *Jarid2* reprograms the chromatin accessibility landscape in pituitary progenitors promoting exit of stemness and initiation of lineage differentiation.

### Multi-modal hits imply downstream mechanisms of *Jarid2*

To further probe the downstream mechanisms of *Jarid2*, we compared genes with changed gene activity to DEGs identified by SnRNAseq within the same cluster. In conformity with previous IHC at E16.5 (Fig. 4J-N), *Pax7* has reduced mRNA expression and gene activity in *Sox2* Prog (Fig. 7G). It is worth mentioning that *Tbx19* is not changed in any of the three modalities measured (Fig. S4A-C). Another TF involved in POMC lineage differentiation, *Neurod1*, is also not changed in corticotrope or *Sox2* Prog in any of the modalities we examined (Fig. S4D-F). The binding activity of PAX7 is also not changed (Fig. S4I). These results further confirm that *Jarid2* is regulating embryonic POMC lineage differentiation at the *Pax7* progenitor level, rather than regulating the function of PAX7. Additionally, we identified 4 multi-modal hits that have upregulated RNA expression and chromatin accessibility: *Notch2* in *Pou1f1* Prog, *Uty*, *Sulf1*, and *Lgr4* in *Pou1f1* Com, and *Meis2* in *Sox2* Prog.

Based on changes in pituitary stem cells, we decided to focus on *Meis2* (Fig. 8A,B). *Meis2* is expressed in *Prop1* Prog and PP in *Jarid2^Fl/Fl^*control pituitaries, but its expression expanded to *Prop1-Pou1f1*, *Pou1f1* Prog, and *Sox2* Prog in *Jarid2^PitKO/PitKO^* (Fig. 8A), resulting in significant increases in mRNA levels. The same pattern is also reflected in the chromatin accessibility level, with an upregulation in *Sox2* Prog, PSC, *Prop1* Prog, *Prop1-Pou1f1*_1, *Pou1f1* Prog, *Pou1f1* Com, and Thyro (Fig. 8B). A closer look at the Tn5 insertion peak pattern of *Meis2* in *Sox2* Prog further unveils widespread increase of chromatin accessibility across the gene body of *Meis2* (Fig. 8C). Previously published Chromatin Immunoprecipitation Sequencing (ChIP-seq) on mouse embryonic stem cells identified binding sites for JARID2 in *Meis2* gene body (Kanellopoulou et al., 2015; Sanulli et al., 2015), shown in Fig. 8C in yellow arrows), with the two closer to the promoter co-occupied by H3K27me3 (Kanellopoulou et al., 2015). We validated the upregulation of *Meis2* in *Jarid2^PitKO/PitKO^* by RT-qPCR in p0 pituitaries (Fig. 8D), further confirming *Jarid2* regulates *Meis2* mRNA level in the pituitary at two different ages. The chromatin accessibility associated with the MEIS2 binding motif is not altered (Fig. S4J), indicating that the change is occurring at the *Meis2* transcriptional level.

**Fig. 8.**
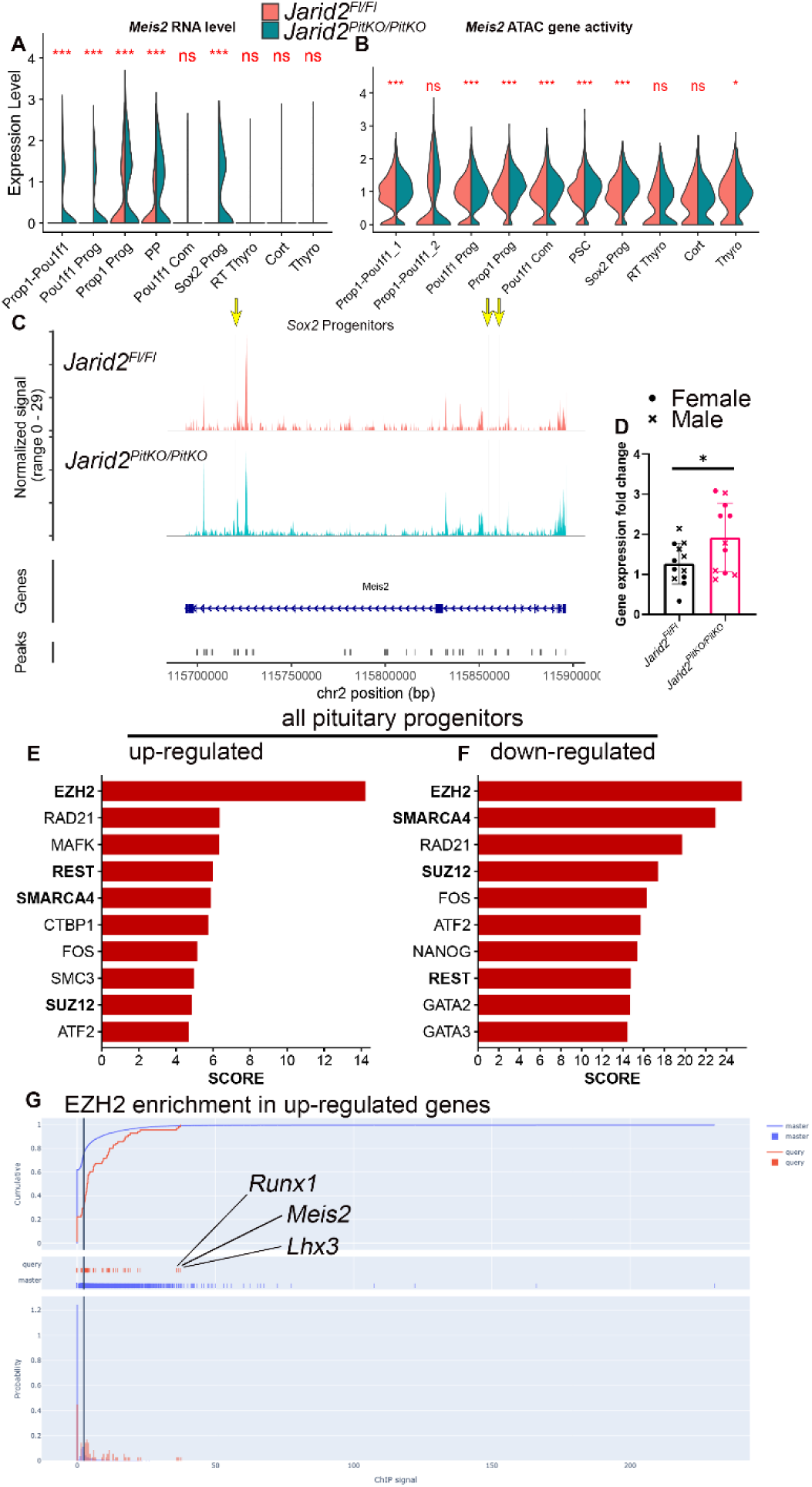
Identification of potential downstream effectors of *Jarid2* in pituitary progenitors. (A,B) Violin plot of the RNA level (A) and gene activity level (B) of *Meis2* in all pituitary clusters. (C) Chromatin accessibility peak map at the *Meis2* locus in *Jarid2^Fl/Fl^* and *Jarid2^PitKO/PitKO^*. Yellow arrows indicate JARID2 binding sites previously published. (D) Quantitative reverse transcription polymerase chain reaction (qRT-PCR) comparing *Meis2* mRNA levels in *Jarid2^Fl/Fl^* and *Jarid2^PitKO/PitKO^* p0 pituitaries shows a significant increase. “*” indicates P < 0.05. “●” indicates female. “X” indicates male. (E,F) Box plot showing MAGIC prediction of transcription factors and cofactors driving the up-regulated (E) and down-regulated (F) genes in all pituitary progenitors. Top 10 TFs ranked by their scores are displayed. (G) MAGIC enrichment distribution of EZH2. P = 9.45*10^-12^. *Lhx3*, *Meis2*, and *Runx1* are the top three hits with highest score.

To further mine changes in TF regulatory networks in pituitary progenitors from the snRNAseq, we utilized Mining Algorithm for Genetic Controllers (MAGIC) to infer enrichment of TFs and cofactors in the gene body and flanking regions of DEGs identified by Wilcoxon’s rank sum test in Seurat (Roopra, 2020). This algorithm predicts regulators based on published ENCODE ChIP-seq data and can infer co-factors without a DNA binding domain. With this method, we identified that both up-regulated and down-regulated genes are enriched for EZH2 and SUZ12 (Fig. 8E,F), indicating that *Jarid2* is functioning as a PRC2 complex subunit in pituitary progenitors. Top three genes predicted to be regulated by EZH2 are *Lhx3*, *Meis2*, and *Runx1* (Fig. 8G). Since *Lhx3* and *Runx1* are both important for pituitary stem cells (Rizzoti et al., 2023), these results further implicate the potential role of *Meis2* co-regulated by JARID2 and EZH2 in pituitary stem cells. We have also identified the REST complex as a potential regulator of *Jarid2*’s function in pituitary progenitors. The epigenetic co-factor of REST complex CoREST has been shown to regulate pituitary hormone expression (Wang et al., 2007). Additionally, the core catalytic subunit of the chromatin remodeling complex Brg/Brm-associated factor (BAF, the mammal homolog of Switch/Sucrose Non-Fermentable, SWI/SNF complex), SMARCA4, is also among the top hits of *Jarid2* downstream regulators. Mutations in multiple subunits of the BAF complex, including *Arid1a*, *Arid1b*, and *Dpf2*, have been associated with CH in patients (Martinez-Mayer et al., 2024b; McGlacken-Byrne et al., 2022). Future study is required to determine the role of *Meis2* and BAF complex in pituitary development.

## DISCUSSION

Embryonic pituitary development requires sophisticated coordination of extrinsic signaling pathways and intrinsic TF activity (Edwards and Raetzman, 2018). During these modulations, chromatin landscapes undergo dynamic reprogramming from pituitary stem cells to hormone secreting cells (Drouin, 2016). However, little is known about how chromatin landscapes in pituitary stem cells are reprogrammed to differentiate into hormone secreting cells in vivo. PRC2 and its accessory subunit JARID2 are crucial regulators of embryonic stem cell differentiation and lineage commitment (reviewed in (Loh and Veenstra, 2022)). There have been extensive studies on how they mediate gene silencing and thereby control development in other tissues (Piunti and Shilatifard, 2021). The core catalytic subunit of PRC2, EZH2, has been shown to be highly expressed in pituitary adenoma and regulate pituitary cell line proliferation (Cai et al., 2023; Schult et al., 2015). Nonetheless, how JARID2 controls pituitary development remains unknown. The current study is the first to show how mutations in PRC2 complex subunit *Jarid2* would affect embryonic pituitary stem cell maintenance and cell fate determination.

Our first important finding is that *Jarid2* regulates the stemness of embryonic pituitary stem cells. We characterized the role of *Jarid2* in pituitary stem cells at two levels. First, *Jarid2* regulates the number of embryonic pituitary stem cells. The number of pituitary stem cells is under strict regulation of intricate signaling pathways, including WNT, Notch, and Hippo (Pérez Millán et al., 2024). As pituitary stem cells differentiate, they will exit the niche around Rathke’s pouch and relocate ventrally, accompanied by the exit of cell cycle (Bilodeau et al., 2009; Davis et al., 2013). Our results showed that both *Jarid2^mut/mut^*and *Jarid2^PitKO/PitKO^* lead to mild pituitary stem cell hyperplasia. In *Jarid2^PitKO/PitKO^*, we identified that *Jarid2* loss affects both cleft and parenchyma stem cells at E16.5. To summarize, our study shows that *Jarid2* normally restricts the number of embryonic pituitary stem cells. This is similar to its role in hindbrain neural progenitors (Takahashi et al., 2007) and cardiac cells (Toyoda et al., 2003).

Second, we also demonstrated that *Jarid2* regulates pituitary stem cells at the transcriptional and chromosomal level through sn multiomics. We identified changes in pituitary progenitor mRNA expression profiles, finding genes enriched in neuron development, axon guidance, and neuron migration. Among these genes, *Slit2* and *Nrg1* are recently implicated to be enriched in pituitary stem cells (Kövér et al., 2026). Human mutations of *SEMA3A* and *SLIT2* have both been linked to pituitary defects in patients (Brauner et al., 2020; Hu and Sun, 2019). Additionally, using pseudotime trajectory analysis, we observed a shift in the pseudotime progression in *Jarid2^PitKO/PitKO^* progenitors, suggesting a change in stem cell dynamics induced by *Jarid2* loss. Similar roles of *Jarid2* in regulating the temporal dynamics of progenitors have also been determined in retinal progenitors, where loss of *Jarid2* leads to a retention in early cell types (Zhang et al., 2023). We further explored changes happening in the chromatin accessibility level in pituitary stem cells and identified increased openness in regions containing SIX and TEAD family motifs. SIX family TFs have been implicated to control pituitary stem cells (Gaston-Massuet et al., 2008; Li et al., 2002). Extensive studies have also been conducted on the role of TEAD family TFs and their upstream regulator hippo pathway to show their essential roles in regulating pituitary stem cell properties (Lodge et al., 2019). Our results unveil a novel epigenetic layer in pituitary stem cell control mediated by *Jarid2*.

Our study also establishes that *Jarid2* regulates pituitary stem cell differentiation to the POMC lineage. The function of chromatin remodeling in POMC lineage differentiation is the most well-defined among the three pituitary lineages. After exiting the cell cycle, the POMC lineage initiates with the fate selection of pioneer TF *Pax7* (Drouin, 2022). With *Pax7* expression, a group of melanotrope-specific chromatin regions will open, followed by binding of *Tbx19* (Mayran et al., 2018; Mayran et al., 2019). The function of *Pax7* has been reported to coordinate with DNA methylation and methylation at lysine 4 of histone H3 (H3K4me1) (Harris et al., 2025). *Neurod1* has also been indicated to contribute to corticotrope differentiation in a *Tbx19*-independent manner (Lamolet et al., 2004). In our study, we first discovered a reduction in AL POMC+ cell in both *Jarid2^mut/mut^* and *Jarid2^PitKO/PitKO^*at E14.5, establishing a strong correlation between *Jarid2* and corticotrope cell number. Furthermore, we identified that POMC+ melanotrope and PAX7+ IL progenitor cell numbers are also reduced, suggesting that *Jarid2*’s regulatory role may be to allow the transcription of *Pax7*. On the molecular level, we confirm reduction in both *Pax7* RNA level and chromatin accessibility in *Sox2* Prog. We observe no significant change in *Tbx19* or *Neurod1* in RNA or chromatin accessibility level. Therefore, our results show *Jarid2* regulates POMC cell differentiation potentially through regulating *Pax7*, independently of *Tbx19* or *Neurod1*. We did not identify any changes with *Pou1f1* lineage or *Nr5a1* lineage, suggesting a *Pomc* lineage specific effect. Our finding determines the regulatory role of *Jarid2* in pituitary cell fate determination, further amplifying the epigenetic mechanism underlying POMC lineage differentiation.

Another important contribution of our study is the first snATACseq of the embryonic pituitary. TF hallmarks of pituitary development have been well-established (Edwards and Raetzman, 2018). However, the underlying chromatin landscape during development still needs to be characterized. Studies have endeavored to determine the chromatin landscape of mouse pituitary under different physiological conditions (Ruf-Zamojski et al., 2021; Ruf-Zamojski et al., 2023). These studies focus exclusively on 7-11 week-old mice (Kövér et al., 2026). The chromatin landscape of postmortem human pituitary from pediatric to aged timepoints has also been determined (Zhang et al., 2022). Nonetheless, there is still a research gap in the chromatin landscape during embryonic development. Here we publish the first snATACseq conducted at E14.5, when pituitary lineage commitment commences and pituitary stem cells are active. We found that most pituitary clusters have distinct chromatin accessibility patterns that correlates with the snRNAseq expression, except for the proliferating progenitors which may suggest that proliferation program happens in the transcription level rather than chromatin accessibility level. We have also identified new subclusters of the intermediate progenitors expressing both *Pou1f1* and *Prop1*. Future study is still required to understand the characteristics of these progenitors and reproducible chromatin landscape of the embryonic pituitary.

We also uncovered the spatial distribution pattern of H3K27me3 during pituitary embryonic development. We identified that the bulk level of H3K27me3 increases as cells differentiate and migrate ventrally into the parenchyma of anterior pituitary from E10.5 to E14.5, eventually becoming ubiquitous at E18.5. We are the first study to report the dynamic spatiotemporal pattern of H3K27me3 during embryonic pituitary development. Previous studies in pituitary cell lines established the importance of H3K27me3 for modulating hormone secretion. In the gonadotrope lineage, H3K27me3 levels have been shown to be enriched in the proximal regions and correlate with the expression levels of *Lhb*, *Cga*, and *Gnrhr* (Pnueli et al., 2015; Xie et al., 2017; Yosefzon et al., 2017). For the *Pou1f1* lineage, Daly et al. (2021) previously showed lower H3K27me3 levels correlate with higher chromatin accessibilities and higher RNA levels in *Isl1*, *Gli3* and *Rxrg* locus of the *Pou1f1* progenitor cell line GHF-T1 and the thyrotrope cell line TαT1. In corticotropes, *Ezh2* has been shown to be important for regulating the proliferation of AtT20 corticotrope cell line (Schult et al., 2015). Although we do not observe changes in bulk H3K27me3 level under *Jarid2* mutations, EZH2 and SUZ12, two core components of the PRC2 complex, are predicted to be the top co-regulators of JARID2 from up-regulated and down-regulated genes of pituitary progenitors identified by snRNAseq. This is consistent with the ample studies that characterized JARID2’s role as both agonist and antagonist of the PRC2 complex to maintain a reasonable level of H3K27me3 in a gene-specific manner (Agius et al., 2026; Landeira et al., 2010; Li et al., 2010; Pasini et al., 2010; Sanulli et al., 2015). Future studies on the core components of PRC2 *in vivo* are still required to fully understand the role of H3K27me3 during pituitary embryonic development. We showed that *Jarid2* mutation can cause an up-regulation of *Meis2* RNA levels and chromatin accessibilities specifically in *Sox2* Prog. And we confirmed the upregulation of *Meis2* by q-RT-PCR in p0 mouse pituitaries. MEIS2 is a transcription factor in the MEINOX family identified as cofactors of homeobox (HOX) proteins (Schulte and Geerts, 2019). The critical roles of *Meis2* have been implicated in cranial and cardiac neural crest (Machon et al., 2015), mandibular arch (Fabik et al., 2020), and forehead development (Leung et al., 2022). Overexpression of *Meis2* has been shown to accelerate neurogenic progression in retinal progenitors (Leavey et al., 2025). Since the pituitary also has neural crest contributions (Davis et al., 2016b) and shares similar regulatory programs with retina (Medina-Martinez et al., 2009), it is very likely that *Meis2* also contributes to pituitary development. Moreover, MEIS2 mutations in human patients have been linked to cleft palate, intellectual disability, cranial facial disorder, and developmental delay (Douglas et al., 2018; Giliberti et al., 2020), all of which frequently co-occur with CH. Therefore, our current study establishes a repressive relationship between *Jarid2* and *Meis2*. Further research is still needed to fully elucidate the function of *Meis2* in pituitary development.

### Limitations of this study

Our study has several limitations. First, we focused our efforts primarily on embryonic pituitary stem cells. The pituitary expands postnatally through both stem cell proliferation and proliferation of intermediate progenitors and differentiated cells. It is still unclear how *Jarid2* would function in the postnatal growth of pituitary. Second, due to the small size of the embryonic pituitary, it is hard for us to have direct binding evidence of JARID2 in chromatin complexes, nor could we determine the developmental profile of H3K27me3 by ChIP-seq or CUT&Tag. For the sn multiomics, we pooled pituitaries from six mixed-sex embryos, but they were still one technical replicates since they were sequenced together. Further study on more technical replicate is needed to make reproducible conclusions about the embryonic pituitary chromatin landscape. Finally, we did not do any direct functional analysis to prove that *Meis2* is actually regulating pituitary development. Future studies are still required to address all these shortcomings.

## ACKNOWLEDGEMENTS

This work was supported by National Institutes of Health (NIH) T32HD108075. We are grateful for Dr. Karine Rizzoti for kindly providing *Prop1*-Cre mice. We acknowledge Dr. Avtar Roopra for help with MAGIC analysis. We thank Dr. Sally Camper, Dr. Shannon Davis, Dr. María Inés Perez Millán, Dr. Buffy Ellsworth, and Dr. Leonard Cheung for thoughtful discussions. And we acknowledge Tyler Bell for technical assistance.

## DATA AND RESOURCE AVAILABILITY

Single-nucleus multiomics data in embryonic day 14.5 pituitaries described in this study are available on the National Center for Biotechnology Information Gene Expression Omnibus (Edgar et al., 2002) using accession numbers [Data upload still in progress. We will update the accession numbers once we have them.].

## AUTHOR CONTRIBUTIONS

Z. L. and L. T. R. designed the study, interpreted the data and wrote the manuscript; Z. L., M. L. B., K. E. W., H. B. performed the experiments.

