## Supplementary Figures for "PRC2 complex subunit JARID2 regulates embryonic pituitary stem cell differentiation to the POMC lineage"

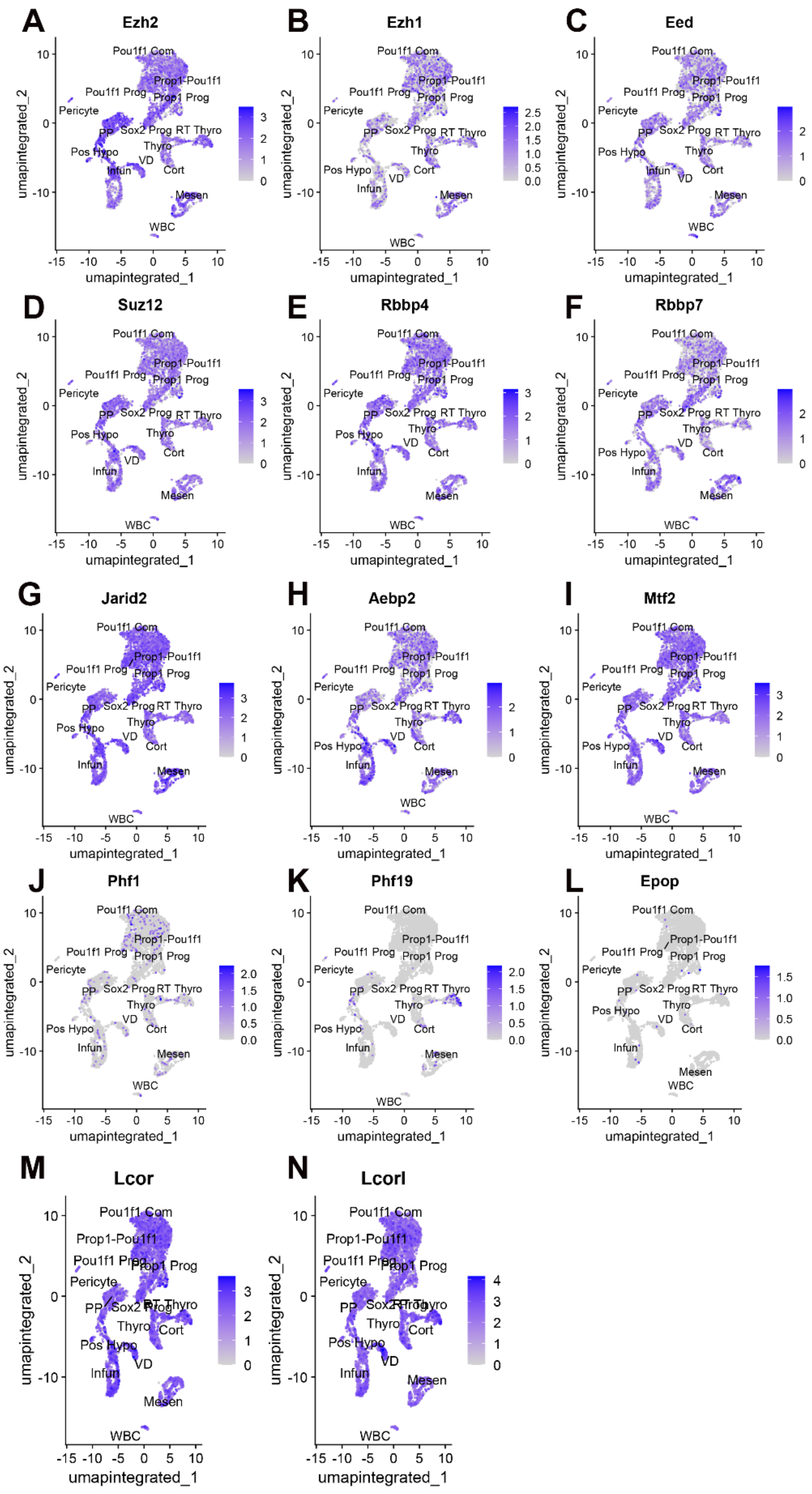

A
Ezh2
B
Ezh1
C
Eed
D
Suz12
E
Rbbp4
F
Rbbp7
G
Jarid2
H
Aebp2
I
Mtf2
J
Phf1
K
Phf19
L
Epop
M
Lcor
N
Lcorl
Pou1f1 Com
Prop1-Pou1f1
Pou1f1 Prog
Prop1 Prog
Pericyte
PP
Sox2 Prog
RT Thyro
Thyro
Pos Hypo
VD
Cort
Infun
Mesen
WBC
umapintegrated_1
umapintegrated_2

**Fig. S1. SnRNAseq in control E14.5 pituitaries characterizes expression patterns of PRC2 core and facultative subunits.** Feature plots of PRC2 core components: *Ezh2* (A), *Ezh1* (B), *Eed* (C), *Suz12* (D), *Rbbp4* (E), *Rbbp7* (F). All core components are ubiquitously expressed. Feature plots of PRC2 facultative subunits: *Jarid2* (G), *Aebp2* (H), *Mtf2* (I), *Phf1* (J), *Phf19* (K), *Epop* (L), *Lcor* (M), *Lcol* (N). Except for *Phf19*, facultative subunits of PRC2 are ubiquitously expressed. *Phf19* is specifically expressed in rostral tip thyrotropes.

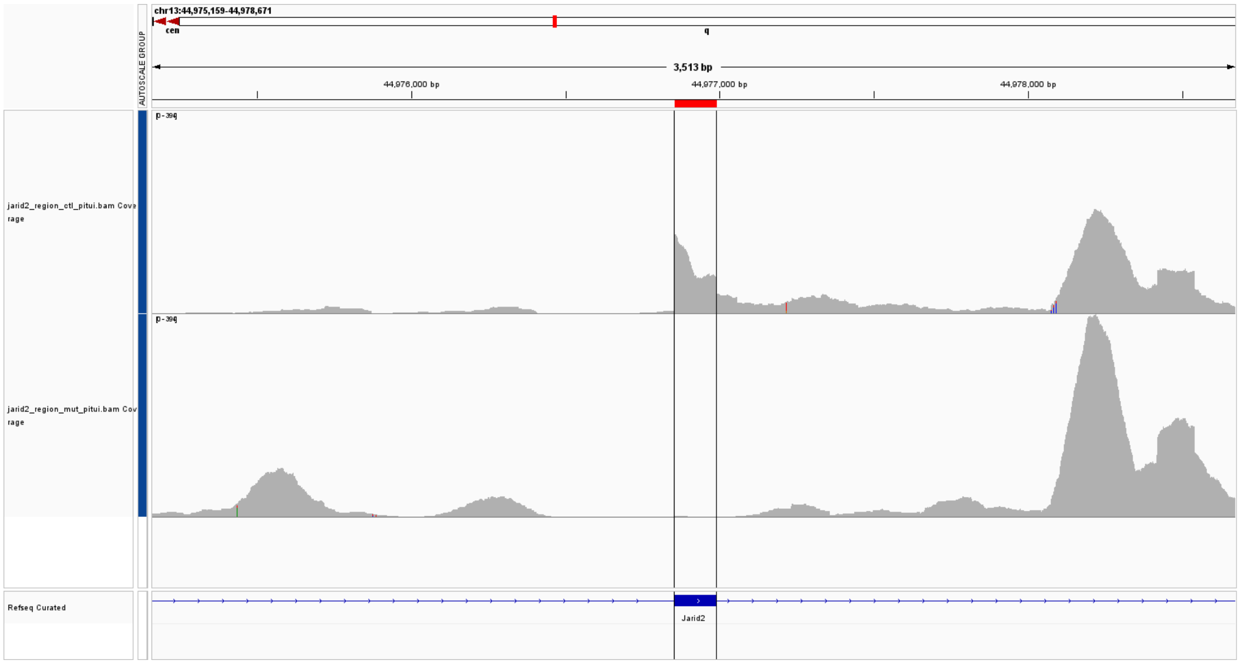


**Fig. S2. SnRNAseq shows deletion of *Jarid2* exon3 specifically in *Jarid2*$^{PitKO/PitKO}$ pituitaries.** Read visualized by IGV 2.19.2.

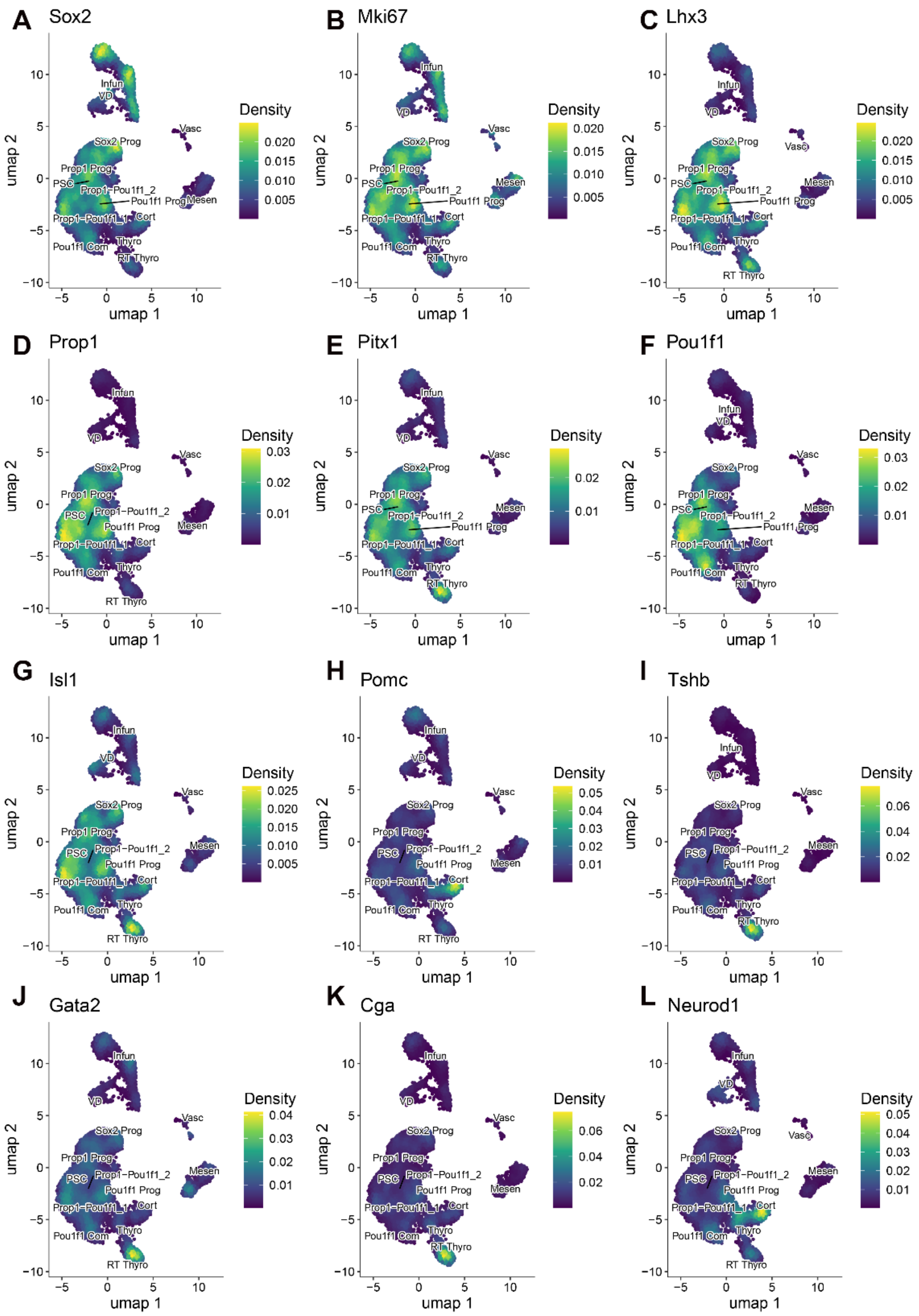


**Fig. S3. Density plot of snATACseq gene activity of marker genes validates the cluster annotation transferred from snRNAseq.** The chromatin accessibility enrichment of *Sox2* (A), *Mki67* (B), *Lhx3* (C), *Prop1* (D), *Pitx1* (E), *Pou1f1* (F), *Isl1* (G), *Pomc* (H), *Tshb* (I), *Gata2* (J), *Cga* (K), and *Neurod1* (L).

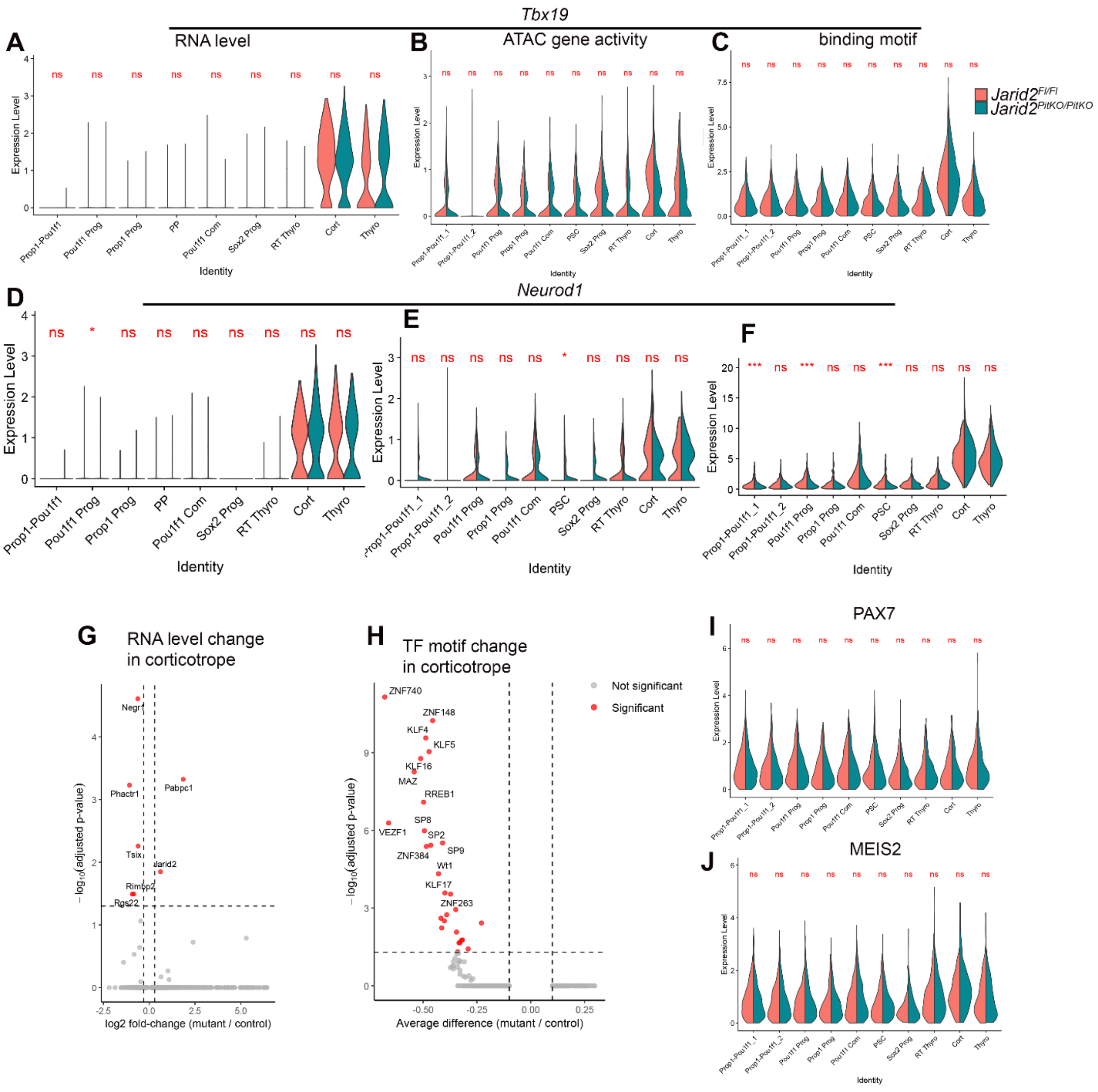


**Fig. S4. Multi modal profiling shows a few changes in corticotropes.** (A-C) No significant change is identified in *Tbx19* mRNA level (A), gene activity (B), or TBX19 binding motif activity (C). (D) Volcano plot showing few genes are changed in corticotrope mRNA levels. (E) Volcano plot showing down regulated TF motif in the corticotrope cluster. (F,G) The binding motif of PAX7 (F) and MEIS2 (G) are not changed. “ns” means not significant.
